# De novo design of functional RNAs through higher-order interactions

**DOI:** 10.64898/2026.09.26.754601

**Authors:** Tongwei Yuan, Dong Wang, Xin-Long Chen, Han-Lin Tao, Chen-Chen Zheng, Xiao-Cong Zhao, Ya-Lan Tan, Xing-Hua Zhang, Zhi-Jie Tan

## Abstract

Designing RNA sequences that reliably adopt functional three-dimensional structures remains a central challenge in RNA engineering because folding depends on cooperative interactions beyond canonical base pairing. Here we present DS3dRNA, an interaction-based framework for de novo RNA sequence design that combines a three-body statistical potential with physics-guided sequence sampling and supports design against multiple conformations. Across the evaluated benchmarks, DS3dRNA outperformed representative RNA inverse-design methods in native-sequence recovery and agreement between predicted and target structures. Energy–sequence-quality analyses further showed that lower design energies generally accompanied higher sequence recovery and macro-averaged F1 scores (MacroF1). Experimentally tested Mango II designs retained high-affinity fluorogenic activity, and five twister ribozyme designs yielded mean endpoint cleavage fractions of 37.7–50.6%, compared with 23.5% for the wild type. These results establish explicit higher-order interaction scoring as a complementary approach to emerging data-driven RNA design methods and provide a framework for designing functional RNAs from experimental or predicted structural ensembles.

## Main

RNA has emerged as a programmable molecular material with broad applications in synthetic biology, therapeutics and biotechnology^1,2,3,4^. Engineered ribozymes, riboswitch-based sensors, fluorescent RNA aptamers and RNA nanostructures illustrate the growing potential of RNA as a functional design platform^1,3,4,5,6^. These applications depend not on sequence or canonical base-pairing patterns alone, but on the ability of RNA molecules to adopt defined three-dimensional (3D) structures and access specific conformational states^4,5,7,8,9,10^. The rational design of RNA sequences that autonomously fold into desired 3D structures therefore represents a central challenge in engineering functional RNAs^11,12,13,14,15^.

Early efforts in 3D RNA design included a Rosetta^16,17^ full-atom framework introduced in 2010 for predicting and redesigning noncanonical RNA motifs^11^. This work explored an energy-based approach to inverse design, in which nucleotide identities are optimized on a prescribed 3D scaffold rather than held fixed for structure prediction. The underlying idea is to search sequence space for nucleotide combinations with favorable estimated interaction energies in the target conformation^11,18,19^. Extending such approaches from individual motifs to more general RNA folds, however, remains challenging. RNA exhibits a complex and dynamic sequence–structure–function relationship: distinct sequences can adopt similar folds, whereas a single sequence may populate multiple conformational states^5,7,12,20,21^. Additional challenges include sparse and structurally biased experimental data^22,23,24,25^, the cost of functional validation^9,21,26,27^, limitations in RNA 3D structure prediction^28,29,30,31,32^, and the difficulty of developing energy functions that adequately capture the cooperative interactions and noncanonical contacts underlying tertiary folding^17,18,19^. These factors complicate the development of broadly applicable energy-based methods for de novo RNA design.

Recent advances in artificial intelligence have begun to reshape this landscape. Deep-learning methods for RNA structure prediction, including AlphaFold3^31^, have substantially expanded the ability to generate and evaluate candidate 3D models^28,29,30,33,34,35,36^, while protein inverse-folding systems such as ProteinMPNN^37^ and LigandMPNN^38^ have demonstrated the effectiveness of message-passing neural networks^39^ for generating sequences conditioned on 3D molecular structure^40^. These developments have stimulated a rapidly growing class of 3D RNA inverse-design methods^13,14,41,42,43,44^. Despite these advances, a distinction remains between learning sequences that are statistically compatible with a target backbone and explicitly optimizing the physical interactions that stabilize that backbone^11,16,17,18,19,45^. Most current data-driven approaches learn conditional sequence distributions from experimentally observed or computationally derived RNA structures, with native sequence recovery or related likelihood-based objectives providing a major source of supervision^13,14,42,44^. Such objectives capture sequence–structure relationships represented in the training data, but they do not directly quantify whether a designed sequence preferentially stabilizes the target conformation relative to alternative structural states^7,11,14,45,46^. This distinction is particularly important for RNA, whose tertiary structures emerge from cooperative networks of canonical and noncanonical contacts, base stacking, long-range interactions and conformational coupling^5,7,18,19,20^. De novo 3D RNA design therefore requires not only identifying sequences compatible with local structural environments, but also capturing the higher-order interaction networks that collectively stabilize a target fold^17,18,19^.

To address this challenge, we developed DS3dRNA, a rational de novo 3D RNA design framework driven by higher-order intra-RNA interactions. DS3dRNA incorporates a three-body statistical potential^19^ that explicitly describes higher-order structural correlations, together with local thermodynamic constraints^47,48^, to explore sequence space for candidates predicted to stabilize a prescribed coarse-grained RNA 3D scaffold^18,49,50^. Unlike approaches that rely on learning a direct structure-to-sequence mapping from large training sets^13,14,42^, DS3dRNA evaluates candidate sequences through an explicit interaction model, providing a complementary physics-inspired route to RNA inverse design. We systematically evaluated DS3dRNA against representative 3D RNA design methods, including RhoDesign^13^, gRNAde^14^, R3Design^41^, RiboDiffusion^42^, RIdiffusion^43^ and AlignIF^44^. To minimize information leakage and provide a stringent assessment of generalization, we constructed challenging benchmark sets subjected to sequence and structural redundancy filtering, and further evaluated targets from CASP15^24^, CASP16^25^ and CASP17^51^. Across these benchmarks, DS3dRNA outperformed these representative methods on both sequence-level and structure-level measures, including native-sequence recovery, MacroF1, and AlphaFold3-based structural consistency assessed by C4′ root-mean-square deviation (RMSD) and TM-score. DS3dRNA also supported multistate RNA design and outperformed the multistate-capable comparator gRNAde under the corresponding benchmark setting. Finally, we experimentally tested DS3dRNA-designed functional RNAs, including twister self-cleaving ribozymes^8,52^ and Mango II fluorescent aptamers^5,9,53^.

## Results

### Higher-order interactions guide sequence design on RNA scaffolds

DS3dRNA formulates inverse design as sequence optimization on a prescribed 3D scaffold (Fig. 1). Each nucleotide is represented by P, C4′ and N1/N9 sites. Spatial triplets are scored with a knowledge-based three-body potential derived from 3,260 structures curated from RNAsolo 2.0^19,22^, and Template-centered alignment groups supply weighted interaction counts for two resolutions of the potential: Rough supports broad sequence exploration, and Fine ranks finalists (Fig. 1a). Local nearest-neighbor screening^47,48^ complements tertiary scoring; optional fixed-position and base-pair constraints restrict allowed mutations. A target may comprise one coordinate set, several residue-matched conformers, or mapped structural fragments, extending established RNA assembly concepts^17,50,54^.

**Fig. 1.**
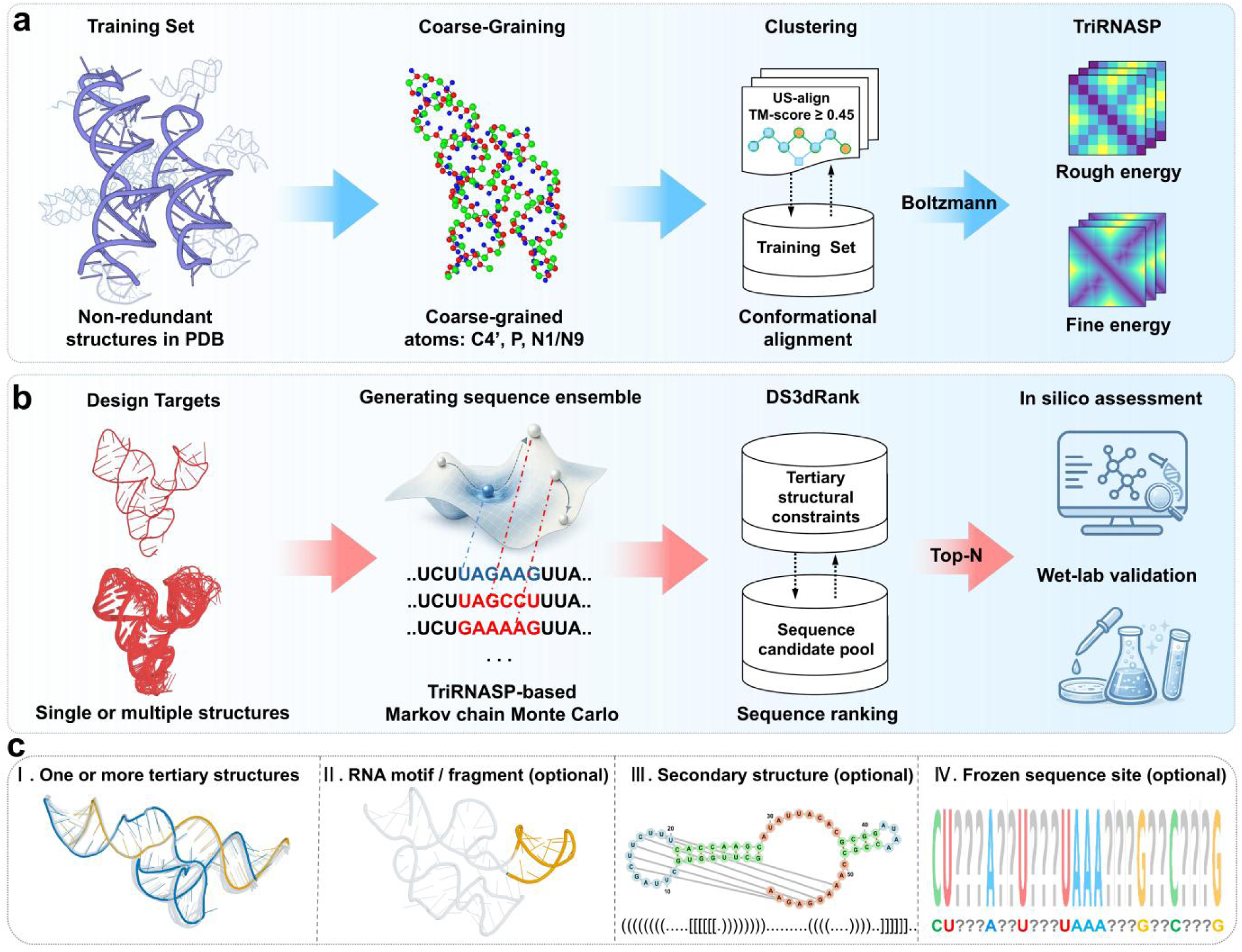
Overview of the DS3dRNA framework for structure-guided RNA sequence design. **a**, Construction of the three-body statistical potential. Experimentally determined RNA structures are represented by P, C4′ and N1/N9 sites and grouped by structural alignment to estimate interaction statistics. Template-centered alignment groups use a C4′-based US-align TM-score threshold of 0.45. Boltzmann-law yields coarse (Rough) and fine (Fine) energy functions for sequence sampling and evaluation. b, Design workflow. A single scaffold or a set of residue-matched conformations guides TriRNASP-based Markov chain Monte Carlo sampling. DS3dRank evaluates candidate sequences against the structural constraints, and selected candidates are assessed by structure prediction or functional experiments. c, Supported inputs and optional constraints: one or more tertiary structures, mapped RNA motifs or fragments, secondary-structure constraints and fixed nucleotide identities at selected positions.

The workflow first converts target coordinates into a three-site scaffold and obtains pairing constraints from an external annotation or from coarse-grained geometry (Fig. 1b,c). We evaluated scaffold-only DS3dRNA, DS3dRNA_auto with internally inferred base pairs, and DS3dRNA_SS with externally supplied secondary structure. Here, geometric inference builds on the premise of coarse-grained RNA contact modeling^18,55^, while external pairing can be annotated with DSSR or MC-Annotate^56,57^. At each sampling step, 320 sequences are scored with Rough; energy-weighted proposals reduce these to 32 candidates, thermodynamic screening to five, and Fine scoring to one Metropolis proposal^58^ (Supplementary Fig. S1). Adaptive mutation and annealing continue over 10,000 steps, after which retained sequences yield a constrained per-run output sequence and an independently recorded minimum-Fine-energy candidate (_Emin). Each target contributed 100 independent designs. Sequence metrics were summarized within targets according to the output type before dataset aggregation, whereas structural evaluation used consensus candidates and DS3dRNA _Emin candidates separately.

### DS3dRNA outperforms representative methods in native-sequence recovery

We compared native-sequence recovery and macro-averaged F1 score (MacroF1) on T83, a redundancy-filtered set of 83 RNA structures. The comparison included the publicly released RhoDesign^13^, gRNAde^14^, R3Design^41^, RiboDiffusion^42^, RIdiffusion^43^ and AlignIF^44^. The implementations and their applicable input configurations (Table 1). Every method used the same targets, and each target contributed equally to the dataset means (Fig. 2a,b; Supplementary Table S2 and Supplementary Data 1).

**Fig. 2.**
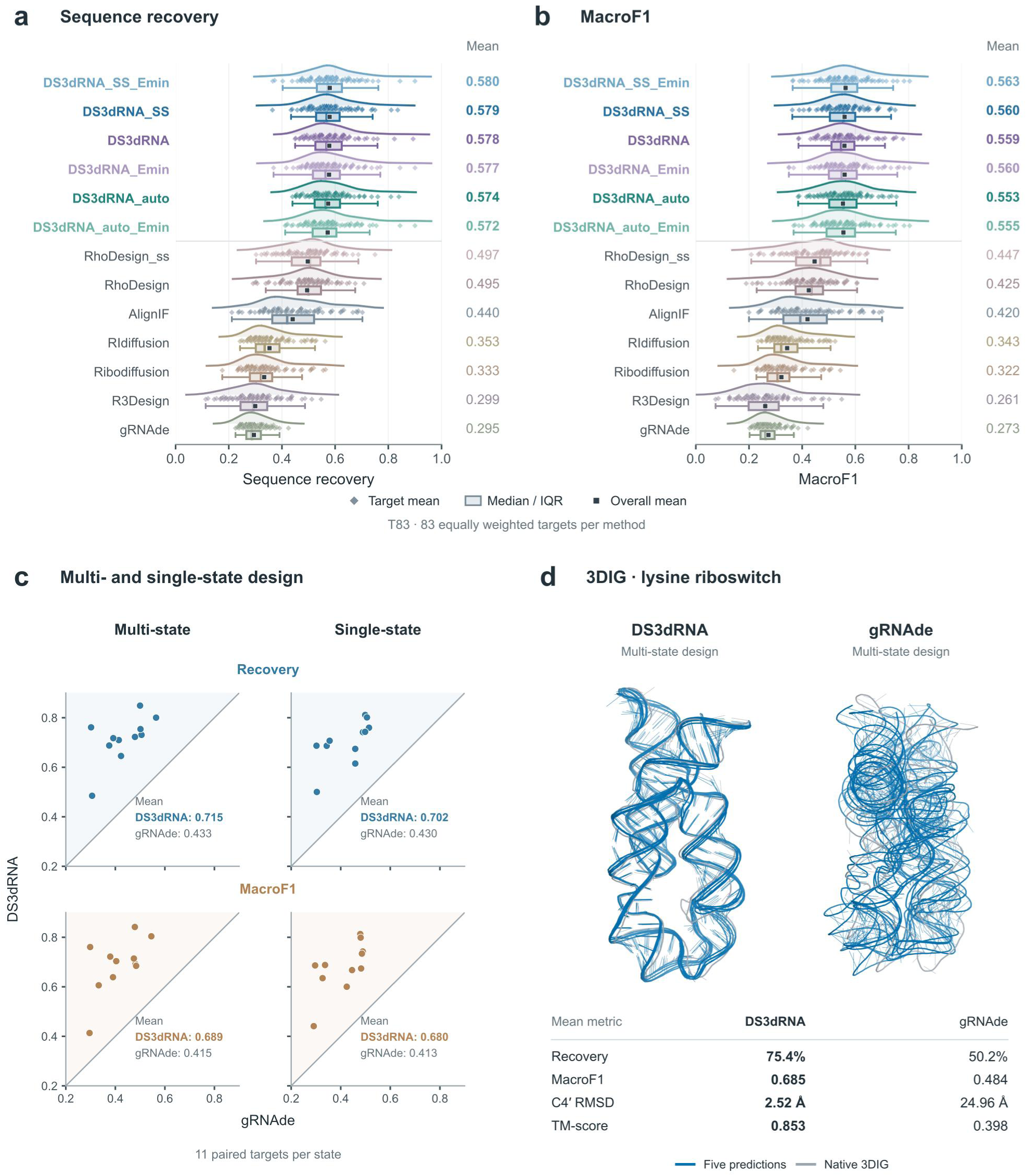
Sequence recovery and multistate RNA design. **a,b**, Sequence recovery (a) and MacroF1 (b) on T83 for 13 configurations and outputs, including all six DS3dRNA entries (83 equally weighted targets per entry). Diamonds denote target-level scores: retained-run means for ordinary outputs and single-candidate scores for _Emin outputs. Boxes span the interquartile range, central lines mark medians and whiskers extend to the most extreme observations within 1.5 times the interquartile range; all observations are shown. Squares and labels give arithmetic means across targets. Curves are boundary-reflected Gaussian kernel density estimates with a common bandwidth within each panel and peak-normalized heights. c, Paired comparisons of DS3dRNA and gRNAde under multi-state and single-state design on T11 (11 targets i.e., ensemble). Points represent targets; rows show recovery and MacroF1, and diagonals indicate equality. Labels give target-level means. d, Structural example for the 174-nt lysine riboswitch (PDB 3DIG). Five AlphaFold3 predictions per multi-state design (blue) are superposed on the experimental reference (grey). C4′ RMSDs are averaged over five predictions, each least-squares fitted to reference chain A using all 174 position-matched C4′ atoms. Recovery, MacroF1 and TM-score summaries are shown below. Length-dependent comparisons are provided in the Supplementary Information.

**Table 1.**
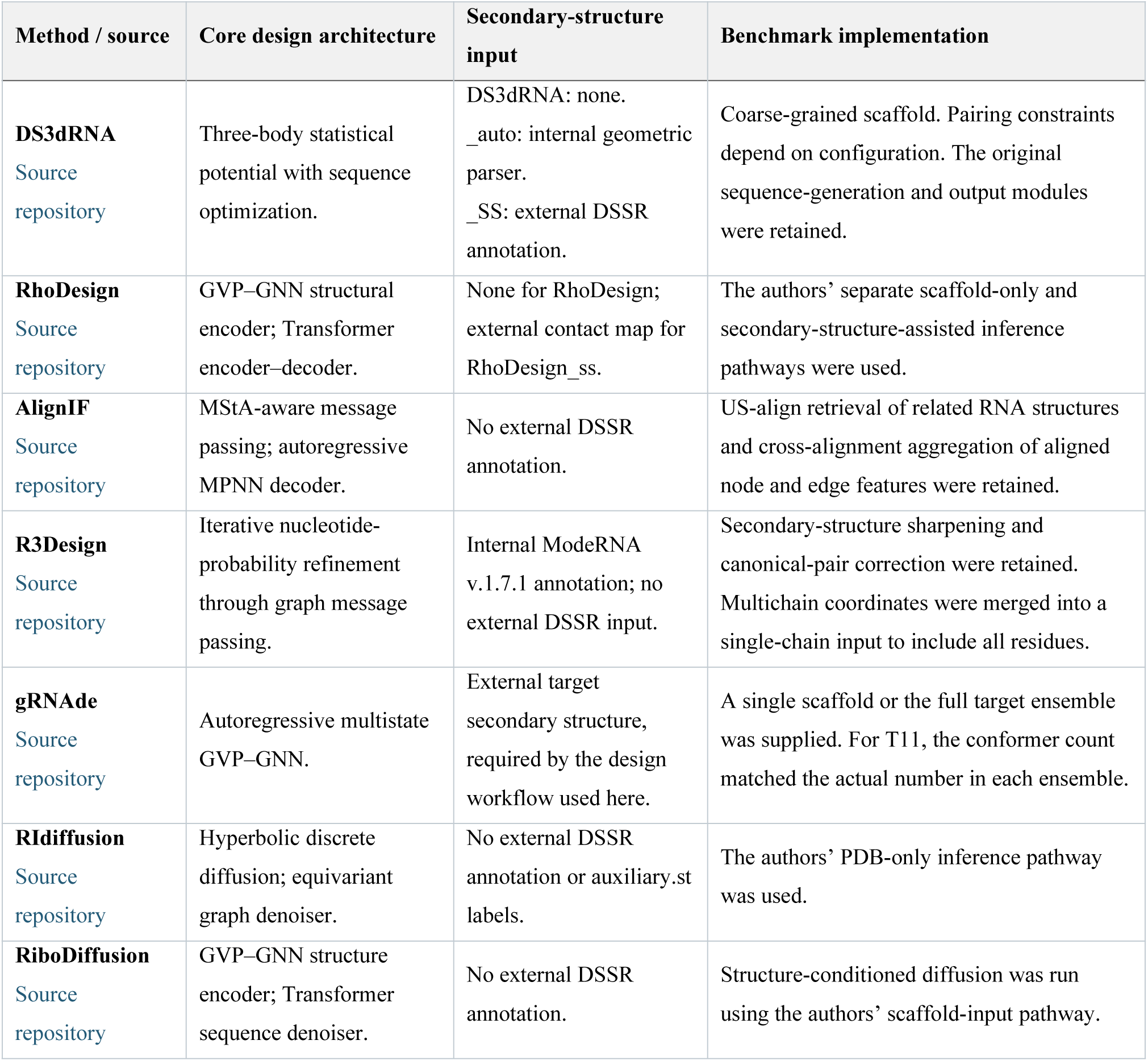
RNA design methods, structural inputs and benchmark inference configurations. The table summarizes the released implementations used for comparison, their treatment of secondary-structure information and the inference procedures retained in this study. DS3dRNA, DS3dRNA_auto and DS3dRNA_SS denote scaffold-only, internally inferred pairing and externally supplied pairing configurations, respectively. The _Emin suffix denotes the additional minimum-Fine-energy output evaluated separately from the consensus candidate. GVP–GNN, geometric vector perceptron graph neural network; MPNN, message-passing neural network.

DS3dRNA_SS reached mean recovery of 0.579 and MacroF1 of 0.560, compared with 0.497 and 0.447 for RhoDesign_ss, the strongest performance of non-DS3dRNA methods on both metrics. Scaffold-only DS3dRNA reached 0.578 and 0.559, and DS3dRNA_auto reached 0.574 and 0.553; their _Emin outputs ranged from 0.572 to 0.580 in recovery and from 0.555 to 0.563 in MacroF1. Among the other methods, RhoDesign scored 0.495/0.425, AlignIF 0.440/0.420, RIdiffusion 0.353/0.343, RiboDiffusion 0.333/0.322, R3Design 0.299/0.261 and gRNAde 0.295/0.273 (recovery/MacroF1), respectively. The DS3dRNA distributions shifted toward higher target-level scores, whereas several comparators showed a broader low-score tail (Fig. 2a,b). Notably, the DS3dRNA advantage was retained without externally supplied base-pair annotations. Moreover, the length-stratified T83 analysis used intervals of ≤50, 51–100, 101–150, 151–200 and 201–350 nucleotides, containing 12, 33, 28, 6 and 4 targets, respectively (Supplementary Figs. S2 and S3). Across these bins, the DS3dRNA configurations maintained comparatively high mean recovery and MacroF1. In the longest interval, DS3dRNA_SS averaged 0.555/0.551 versus 0.367/0.292 for RhoDesign_ss (recovery/MacroF1). The individual-target plots show a broader low-scoring tail for RhoDesign, particularly among longer targets, while the corresponding DS3dRNA observations remain more concentrated at higher scores.

### Lower design energy is associated with higher sequence quality

We related design energy to sequence quality using retained records from 100 runs per target and configuration across T25 and T83. The 720-nucleotide 7PTK target provides a visual example: the retained-record energy profiles declined rapidly during early recorded steps and then approached a plateau (Supplementary Fig. S5). Pooled energy–recovery Spearman correlation coefficients were −0.910, −0.944 and −0.942 for DS3dRNA, DS3dRNA_auto and DS3dRNA_SS, respectively, and corresponding energy–MacroF1 coefficients were −0.909, −0.943 and −0.941. The complete target-level profile collection is available through the public dataset archive linked in Data availability and Supplementary Note 1. The association extended across targets (Supplementary Fig. S6): mean energy–recovery Spearman coefficients across the three configurations ranged from −0.793 to −0.773 on T25 and −0.674 to −0.622 on T83, and the corresponding energy–MacroF1 means were −0.774 to −0.751 and −0.660 to −0.615. The coefficients were computed from unbinned retained records within each target–configuration combination and then averaged with equal target weights. The consistent direction suggests that the interaction-based energy captures aspects of compatibility with native nucleotide assignments that favor reference-like sequences within the sampled design sequence space.

### Joint design accommodates multiple conformational realizations

We evaluated multistate design on T11, comprising ten solution-NMR ensembles and one 174-nucleotide lysine-riboswitch ensemble, with 168 coordinate models in total. Each ensemble contributed one target-level observation. Multistate DS3dRNA reached mean recovery of 0.715 and MacroF1 of 0.689, compared with 0.433 and 0.415 for multistate gRNAde (Fig. 2c). Joint conditioning increased DS3dRNA recovery from 0.702 to 0.715 and MacroF1 from 0.680 to 0.689. The corresponding gRNAde gains were 0.003 and 0.002. Thus, DS3dRNA maintained substantially higher overall scores and showed a larger incremental benefit from joint rather than single-conformer design.

The lysine riboswitch illustrated how joint design translated to AlphaFold3^31^ predicted structural agreement (Fig. 2d). Multistate sequence recovery was 0.754 for DS3dRNA and 0.502 for gRNAde. For the selected candidates, five-model mean C4′ RMSDs to the 174-nucleotide 3DIG reference were 2.52 and 24.96 Å, respectively, with TM-scores of 0.853 and 0.398. Comparisons with the 12 experimental conformers were performed as one-to-many alignments of this single target. The DS3dRNA candidate therefore combined higher sequence recovery with closer prediction to the experimental riboswitch scaffold.

### DS3dRNA outperforms comparators in predicted structural consistency

To examine target-fold consistency, we analyzed T25, comprising seven RNA-only targets from T83 and 18 targets from CASP15^24^ and CASP16^25^, through predicting 3D structures of the designed sequences by AlphaFold3. Here, AlphaFold3^31^ received candidate sequences without target coordinates or target secondary-structure restraints. Five predicted structures were evaluated for each candidate; structural scores were averaged first within targets and then across the 25 equally weighted targets. Figure 3 presents DS3dRNA_SS_Emin alongside the comparator methods, with all DS3dRNA configurations reported in Supplementary Fig. S4.

**Fig. 3.**
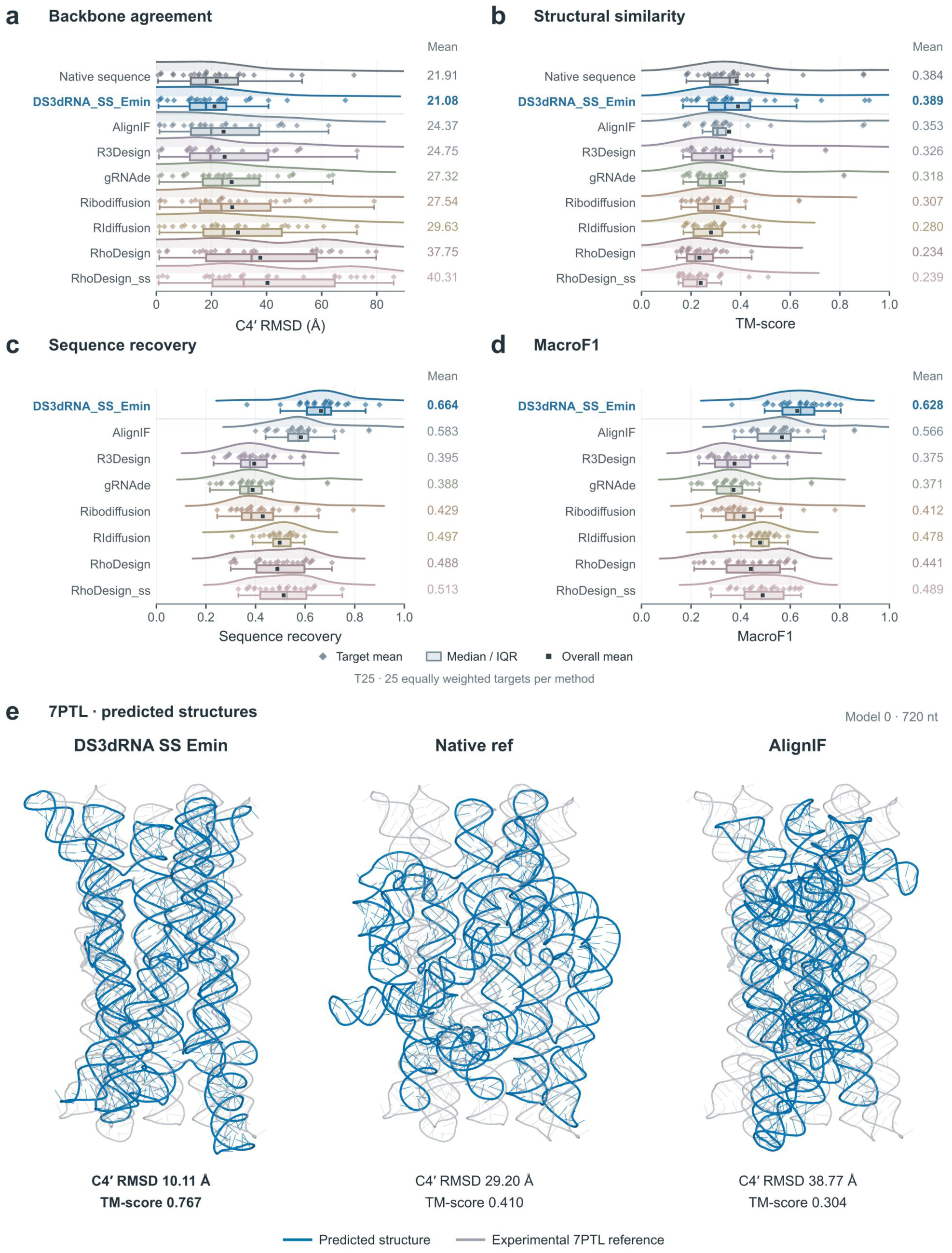
Predicted structural consistency and sequence performance on T25. **a,b**, C4′ RMSD (a) and TM-score (b) for DS3dRNA_SS_Emin, comparator methods and native-sequence predictions. c,d, Sequence recovery (c) and MacroF1 (d) for the design methods; DS3dRNA_SS_Emin uses one minimum-energy candidate per target, whereas comparator values are means across generated sequences. Diamonds represent targets (25 equally weighted targets per method); structural values in a,b average five AlphaFold3 predictions per candidate. Boxes show the interquartile range and median; whiskers extend to the most extreme observations within 1.5 times the interquartile range. All target values are shown. Squares and labels denote means across targets. Curves are boundary-reflected Gaussian kernel density estimates with peak-normalized heights and bandwidths shared with the supplementary all-mode comparison. e, Model-0 predictions for DS3dRNA_SS_Emin, the native sequence and AlignIF for the 720-nt 7PTL target. Predictions are blue and the experimental reference is grey; views share a camera and scale. C4′ RMSD uses least-squares fitting of all 720 position-matched C4′ atoms; TM-score uses the US-align structural correspondence and reference-length normalization. Values in e describe the displayed models, whereas a,b summarize five-model means. All DS3dRNA configurations are compared in the Supplementary Information.

DS3dRNA_SS_Emin achieved a mean C4′ RMSD of 21.08 Å and TM-score of 0.389 on T25, compared with 24.37 Å and 0.353 for AlignIF, the strongest comparator by these structural means metrics (Fig. 3a,b). R3Design, gRNAde, RiboDiffusion and RIdiffusion yielded RMSD/TM-score pairs of 24.75/0.326, 27.32/0.318, 27.54/0.307 and 29.63/0.280; RhoDesign and RhoDesign_ss yielded 37.75/0.234 and 40.31/0.239. For sequence recovery/MacroF1 (Fig. 3c,d), DS3dRNA_SS_Emin scored 0.664/0.628, marginally higher than AlignIF 0.583/0.566; RhoDesign_ss 0.513/0.489; RIdiffusion 0.497/0.478; RhoDesign 0.488/0.441; RiboDiffusion 0.429/0.412; R3Design 0.395/0.375; and gRNAde 0.388/0.371. RhoDesign retained moderate sequence agreement yet showed the largest structural deviations, illustrating that recovery alone cannot substitute for target-fold assessment. It is noted that predictions based on native sequence yielded the averaged metrics of 21.91 Å and 0.384. Full target– and model-level results are provided in Supplementary Table S3 and Supplementary Data 2–3.

Notably, candidate selection could changed the predicted fold for individual targets. On the 720-nucleotide 7PTL target, DS3dRNA_SS_Emin had a five-model mean RMSD of 12.1 Å and TM-score of 0.726, compared with 34.8 Å and 0.322 for its consensus candidate and 35.5 Å and 0.331 for native-sequence predictions. Figure 3e displays fixed-index model 0 to illustrate the contrast: DS3dRNA_SS_Emin yielded C4′ RMSD 10.11 Å and TM-score 0.767, whereas the native-sequence prediction reached the values of 29.20 Å and 0.410 and AlignIF reached 38.77 Å and 0.304. The displayed structures and their labels are model-specific; the benchmark summary uses all five predictions.

Furthermore, we examined the CASP17 P20 kissing-multiloop scaffold 10ZU^12^ as a separate case study (Extended Data Fig. 1). DS3dRNA_auto consensus and _Emin candidates had five-model mean C4′ RMSDs of 9.88 and 9.71 Å and TM-scores of 0.407 and 0.412, respectively, compared with 18.80 Å and 0.223 for the native sequence. AlignIF, RIdiffusion, RhoDesign_ss, RiboDiffusion, gRNAde and R3Design yielded C4′ RMSD/TM-score pairs of 22.19/0.310, 20.22/0.196, 21.42/0.251, 22.20/0.203, 22.61/0.238 and 23.37/0.224, respectively. Here, the automatic configuration inferred pairing from the masked scaffold without external secondary structure. Fixed-index model 0 is shown for each method and the quantitative structural values were averaged over five predictions per candidate.

### Mango II designs retain fluorogenic activity

We tested functional retention in Mango II^5,9^, whose G-quadruplex core contains interactions absent from conventional dot-bracket annotation. From 100 DS3dRNA design trajectories, we selected the three lowest-energy candidates for experimental evaluation and compared them with the optimized design from each RhoDesign mode (with/without secondary-structure) and the wild-type Mango II (Fig. 4 and Supplementary Table S1). No native nucleotide identities were fixed during Mango II redesign, allowing the algorithm to determine nucleotide identities solely from the structural scaffold and interaction potential. Final RNA concentrations were 0–100 nM at a fixed TO1-3PEG-Biotin concentration of 1 nM. For each replicate, fluorescence was expressed as the relative increase above its corresponding zero-RNA baseline.

**Fig. 4.**
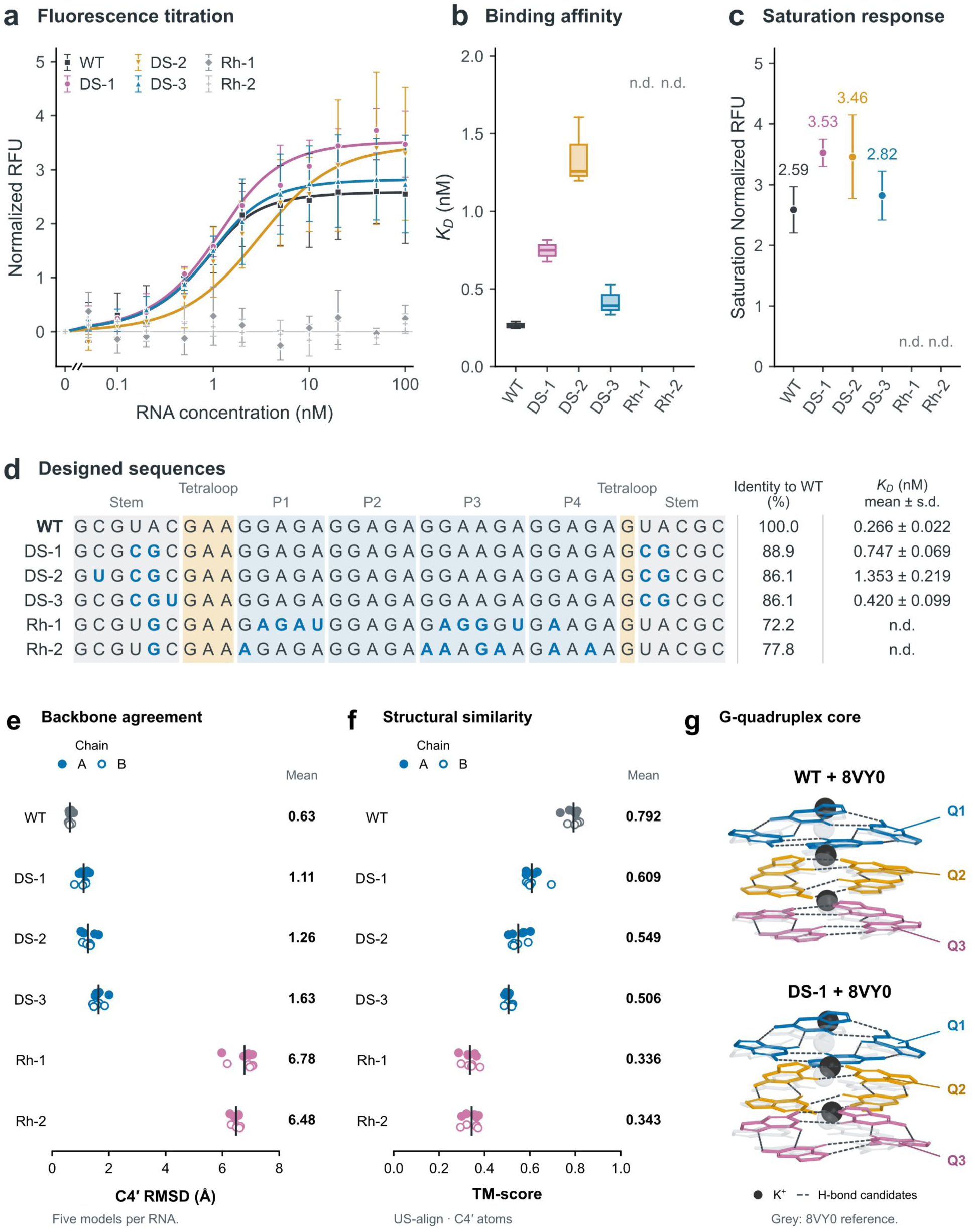
Fluorogenic activity and predicted structures of Mango II designs. **a**, Titration with 1 nM TO1-3PEG-Biotin. Each replicate is expressed as the relative increase above its corresponding zero-RNA baseline. Points show mean ± s.d. (n = 3). Curves are tight-binding fits to mean normalized data, weighted by pointwise s.d., with zero intercept fixed. The concentration axis is linear up to 0.05 nM and logarithmic thereafter. The expanded low-concentration region is shown to visualize the subnanomolar binding transition. b, Apparent KD from separate unweighted replicate fits with measured F0 fixed. Boxes show quartiles and medians; whiskers show the range (n = 3). c, Saturation normalized RFU from a, with covariance-derived standard errors conditional on measured baselines. n.d., not reliably determined. d, Sequences, identity to WT and KD (mean ± s.d., n = 3). Blue marks substitutions. Rh-1/Rh-2 use RhoDesign with/without secondary-structure input. e,f, C4′ RMSD to 8VY0 chain A (e) and C4′ US-align TM-score normalized by the 36-nt reference (f). Filled/open circles distinguish prediction chains A/B; each contains five models. Ticks and labels give means across both chains and five models. g, Predicted WT and DS-1 G-quadruplex cores superposed on grey 8VY0. Q1–Q3 label quartets; spheres denote K^+^ and dashed lines candidate hydrogen bonds.

All three DS3dRNA candidates showed concentration-dependent fluorescence activation (Fig. 4a). Fits to the mean normalized curves gave saturation responses of 3.53 ± 0.23, 3.46 ± 0.69 and 2.82 ± 0.40 for DS-1, DS-2 and DS-3, respectively, compared with 2.59 ± 0.38 for the wild type (fit estimate ± covariance-derived standard error; Fig. 4c). Thus, all three tested DS3dRNA candidates showed robust, saturable fluorescence activation and retained sub– to low-nanomolar apparent affinity, while the two RhoDesign candidates did not produce well-resolved saturating responses in the tested range. Separate fits to three replicate titrations yielded apparent K_D v_alues of 0.747 ± 0.069, 1.353 ± 0.219 and 0.420 ± 0.099 nM for DS-1, DS-2 and DS-3, respectively, compared with 0.266 ± 0.022 nM for wild type (mean ± s.d., n = 3; Fig. 4b,d). All three designs therefore retained sub– to low-nanomolar apparent affinity, within approximately fivefold of the wild type, while producing larger saturation responses. The underlying replicate RFU measurements are provided in Supplementary Table S1.

The three DS3dRNA aptamers retained 86.1–88.9% identity to wild type, with nucleotides 7–30 unchanged in all three; the two RhoDesign candidates had nine and seven substitutions in this region (Fig. 4d). AlphaFold3 predictions of the designed sequences gave mean C4′ RMSDs of 1.11–1.63 Å and TM-scores of 0.506–0.609 against the experimental reference, compared with 6.48–6.78 Å and 0.336–0.343 for the RhoDesign candidates (Fig. 4e,f). For the prediction, two identical copies of the designed RNA sequence were supplied as chains A and B, together with eight K^+^ ions and two TO1-biotin molecules (CCD code EKJ), matching the stoichiometry of PDB 8VY0 and the default settings used seed 1 and five diffusion samples. The above quantitative summaries include both chains across all five predictions, and the G-quadruplex close-ups in Fig. 4g are AlphaFold3 predictions.

### Design from predicted twister ensembles preserves cleavage activity

We next tested design from predicted structural inputs using the bimolecular ES2 twister ribozyme^8^. Five AlphaFold3 models of the wild-type ribozyme–substrate complex were supplied jointly to DS3dRNA. Only the ribozyme strand was redesigned, and the substrate sequence was held fixed. Five low-energy candidates were evaluated against the same Cy3-labeled substrate using the 20-s cleavage assay described in Methods (Fig. 5a–c). Notably, this design tested functional retention when the complete assayed construct was represented by a predicted ensemble rather than an experimentally determined scaffold.

**Fig. 5.**
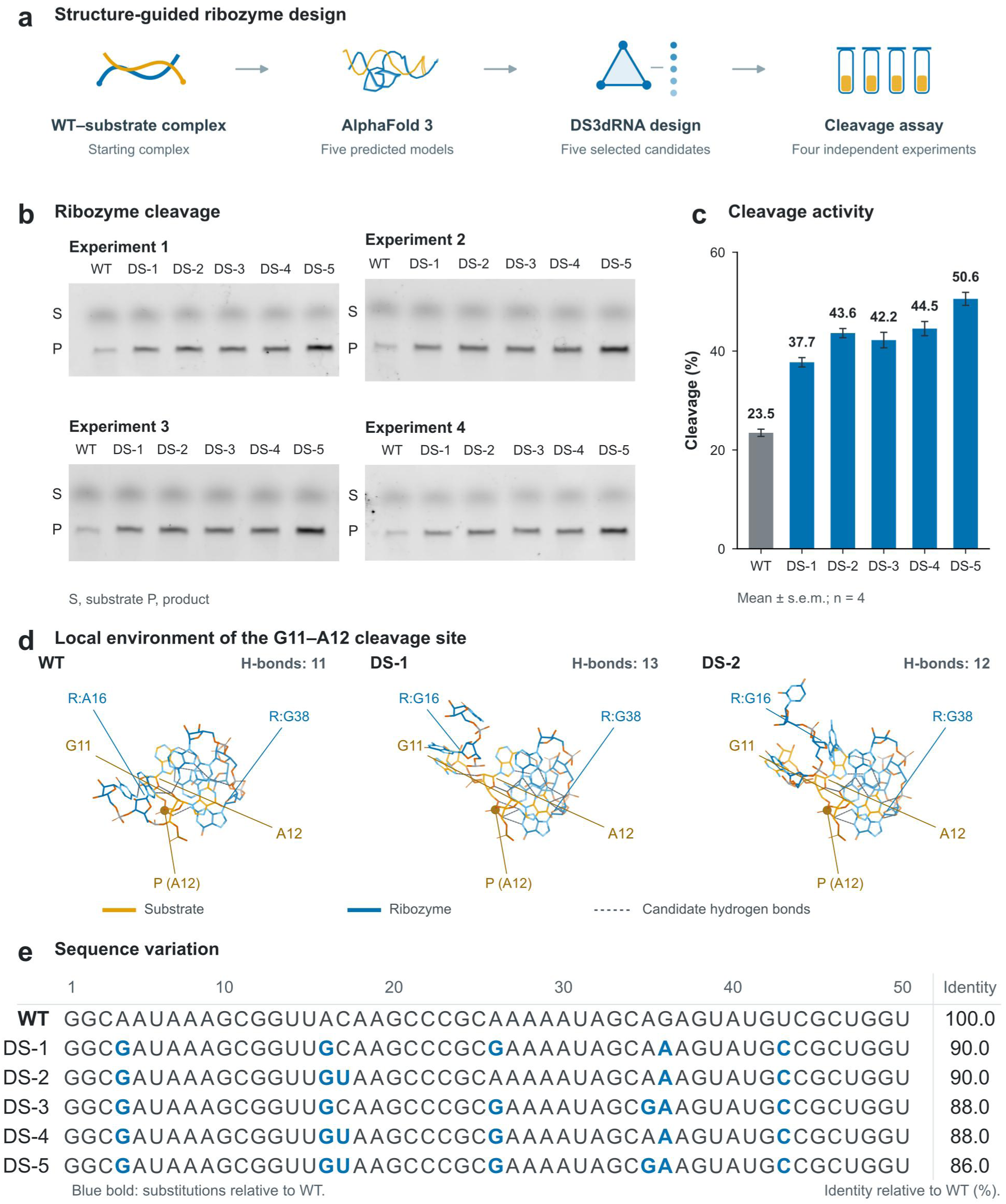
Design from a predicted twister ribozyme ensemble preserves cleavage activity. **a**, Design from five AlphaFold3 models of the WT ribozyme–substrate complex and testing of five DS3dRNA candidates. Only the ribozyme strand was redesigned. b, Denaturing PAGE from four independent experiments. Crops retain six consecutive lanes and substrate (S) and product (P) bands. Experiments 1 and 4 were mirrored horizontally to correct rear-view scan orientation; annotated full-gel images are provided in the Supplementary Information. c, Cleavage at 25 °C after 20 s, shown as mean ± s.e.m. (n = 4); labels give mean percentages. d, Predicted local environments of the substrate G11–A12 cleavage site in WT, DS-1 and DS-2, using model 0 with a common local alignment and camera. Substrate/ribozyme carbons are orange/blue; oxygen/nitrogen are vermilion/light blue. P (A12) marks the A12 phosphorus; R denotes ribozyme residues. Dashed lines show DSSR standard or acceptable hydrogen-bond candidates involving G11 or A12, with heavy-atom distances of 2.5–3.5 Å. The displayed regions contain 11, 13 and 12 candidates, respectively; these predictions do not establish catalytic mechanism. e, The 50-nt ribozyme sequences, with substitutions in bold blue. Identity is the percentage of positions matching WT, excluding the fixed substrate.

At 25 °C, mean substrate cleavage was 23.5% for the wild-type ribozyme and 37.7%, 43.6%, 42.2%, 44.5% and 50.6% for the designed ribozymes DS-1 through DS-5, respectively (n = 4 replicate experiments; Fig. 5c). All five designs retained cleavage activity and showed higher mean 20-s endpoint cleavage than wild type, corresponding to 1.61–2.15-fold increases. The annotated source gels are provided in Supplementary Fig. S8. The five ribozyme sequences retained 86–90% identity to wild type (Fig. 5e). In the model-0 AlphaFold3 views around substrate G11–A12, DSSR and distance criteria identified 11, 13 and 12 hydrogen-bond contacts for wild type, DS-1 and DS-2, respectively (Fig. 5d). The designed structures place the ribozyme-side G38 closer to the local active-site environment, where additional candidate hydrogen bonds appear even as A16 shifts away from its wild-type position. This rearrangement suggests a possible interaction-based route to the higher endpoint cleavage of the tested designs.

## Discussion

DS3dRNA couples an explicit three-body RNA interaction potential^19^ to physics-guided sequence sampling. In silico, DS3dRNA outperformed representative inverse-design methods in native-sequence recovery on T83 and in predicted target-fold consistency on T25. In vitro functional assays further showed that three designed Mango II aptamers retained high-affinity ligand binding and fluorogenic activity, and all five designed twister ribozymes exhibited higher endpoint cleavage than the wild type. Together, these computational benchmarks and functional assays support the utility of higher-order spatial interactions as a source of RNA design information beyond conventional base-pair constraints.

The framework brings tertiary interaction scoring, local thermodynamic screening^47,48^ and optional sequence or pairing constraints into the same search procedure. Similar performance across the three DS3dRNA input configurations on T83 indicates that the target scaffold supplies useful design information even without an external secondary-structure annotation. The negative energy–quality correlations across T25 and T83 further suggest that the scoring function captures aspects of the interaction network associated with reference-like sequence assignments. The example of 7PTL illustrates the effect of candidate selection: minimum-energy ranking identified a sequence with substantially closer predicted target-fold agreement than consensus decoding. Because the score can be evaluated for externally generated sequences, interaction-based ranking could also complement learned sequence generators such as RhoDesign^13^, gRNAde^14^ and AlignIF^44^. Multistate conditioning provides a way to use structural variation during design^14,46^. On T11, DS3dRNA achieved higher recovery and MacroF1 than gRNAde and showed a larger gain when multiple conformers replaced a single structural input. The example of lysine-riboswitch further connected joint design with close predicted agreement to an experimental scaffold. In the twister ribozyme application, the same approach accommodated an ensemble of predicted ribozyme–substrate complexes and produced five active designs. Together, these results support the use of multiple coordinate realizations when a functional RNA cannot be adequately represented by one static structure.

Beyond in silico benchmarking, the two experimental systems provided complementary tests of structure-dependent RNA function. Mango II relies on noncanonical interactions within its G-quadruplex core and a ligand-binding environment that supports fluorescence activation^5,53^. All three DS3dRNA-designed candidates retained subnanomolar to low-nanomolar apparent affinities and showed larger relative increases in fluorescence at saturation than the wild type. The twister ribozyme assay, in turn, probed catalytic function, with all five designs showing higher mean cleavage than the wild type at the measured endpoint. Predicted rearrangements around G38 and A16 suggest a testable structural hypothesis for this increase: additional local contacts could favor catalytically competent configurations despite changes elsewhere in the active-site environment.

In practical RNA engineering, DS3dRNA offers a flexible way to combine structural scaffolds with prior knowledge. Experimentally established residues can be retained while surrounding positions are redesigned, pairing constraints can be supplied or inferred from the scaffold, and multiple conformations or mapped structural fragments can be combined within the same optimization procedure. This flexibility brings motif-preserving redesign, scaffold-guided sequence generation and multistate optimization into one framework. The Mango II and twister experiments demonstrate its utility for producing functional candidates in two distinct RNA systems. Combining these capabilities with broader conformational sampling and quantitative sequence–function landscapes^21,45^ may extend their use in engineering ligand binding, catalysis and molecular switching^27,46^. DS3dRNA therefore provides an adaptable, interaction-based route from available structural and experimental information to testable RNA sequences.

## Methods

### RNA 3D structure representation

#### Coarse-grained representation for RNAs

A key principle in 3D RNA design is that the structural representation of the target scaffold should not directly reveal the nucleotide identity to be designed through nucleotide-specific spatial patterns or atom-type features^14,42^. Therefore, DS3dRNA represents each RNA residue using three coarse-grained (CG) atoms: the backbone atoms P and C4′, and the base-associated atom N1 for pyrimidines or N9 for purines. This CG representation provides a compact residue-level description of RNA tertiary structure while avoiding the full set of nucleotide-specific heavy atoms that encode residue identity.

As illustrated in Fig. 1a, the P (green) and C4′ (red) atoms preserve the essential geometric organization of the RNA backbone, whereas the N1/N9 atom (blue) serves as a residue-level anchor for the spatial placement of the nucleobase. This CG representation is well suited for capturing intrinsic interactions in nucleic acid systems while maintaining computational feasibility^49,50,59^, and similar CG descriptions have been widely adopted in previous approaches to RNA 3D structure modeling, including Vfold^59^, SimRNA^49^, TiRNA^55^, FebRNA^50^, cgRNASP^18^, and TriRNASP^19^. Related residue-level representations have also been employed in recent 3D RNA design models such as gRNAde^14^, RiboDiffusion^42^, RIdiffusion^43^, and AlignIF^44^.

### Construction of higher-order interaction potentials

#### Theory of higher-order interactions in RNA structures

In our previous work, we developed a knowledge-based many-body statistical potential framework (TriRNASP) for predicting RNA interactions^19^. The potential improved RNA 3D structure quality assessment relative to the pairwise potentials and deep-learning-based methods evaluated in that study, supporting the use of higher-order structural correlations. The corresponding multi-body statistical potential energy, Δ*E*, is defined as:

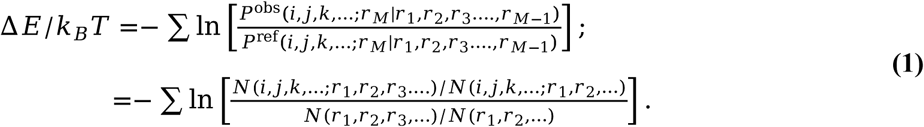

Here, *i, j, k*, … denote atom types, defined by the combination of four nucleotide identities (A, U, C and G) and three bead classes (P, C4′ and N1/N9), and *r*_1_, *r*_2_, *r*_3_, … *r_M_* _d_enote pairwise distances among the *M* CG atoms. The summation is over the atom tuples included in the interaction potential for an RNA structure. *P*^obs^ is the observed conditional probability of *r*_M g_iven *r*_1_, *r*_2_, …, *r*_M−1 i_n native structures, and *P*^ref^ is the corresponding probability in a reference state. In DS3dRNA, the reference distribution was obtained by averaging over nucleotide-specific atom types. Because nucleotide identity is the design variable, this averaging maps the sequence space onto an equal-weight atom-type subspace and therefore corresponds to a maximum-entropy sequence reference; consequently, a completely random observed atom-type distribution gives Δ*E* ≈ 0. For DS3dRNA, we derived the potential at the three-body level, and Eq. (1) reduces to:

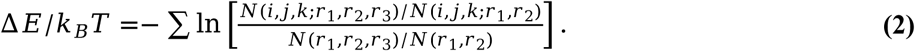

Here, *N*(*variable*) denotes a multidimensional count tensor. In DS3dRNA, distances were discretized into uniform integer-width bins for the Rough and Fine potentials, E_Rough a_nd E_fine (_Fig. 1a), producing spatial tensor dimensions of 7 × 7 × 14 and 18 × 18 × 36, respectively. The cutoffs for r₁ and r₂ were both set to 9.5 Å, sufficient to capture contacts involving the base-anchor N1/N9 bead. The range of r₃ was naturally constrained by the triangle geometry defined by r₁ and r₂, and geometrically inaccessible bins were excluded. An accessible bin with zero counts was assigned the value of the least frequently observed accessible spatial relation within the same atom-type triplet.

#### DS3dRNA training set

The DS3dRNA training set comprised 3,260 RNA structures. To construct this set, we retrieved RNA structures from the RNAsolo 2.0 archive^22^ (accessed 1 December 2025) using the following query settings: structure repository, BGSU; redundancy, all class members; molecule, all; experimental method, all; resolution, ≤4.0 Å; and format, PDBx/mmCIF. Approximately 14,000 entries were retrieved. We retained RNAs containing 16–512 nucleotides and excluded entries containing any homopolymeric chain. Initial redundancy filtering was performed using US-align^60^ (v.20241108), retaining one representative when a structure pair had either a TM-score or an aligned-region sequence identity of approximately 1.000. This yielded 6,968 candidate structures (clean_pool_6968.zip). We further reduced redundancy by retaining a single representative from structure pairs for which both the TM-score and the aligned-region sequence identity were ≥0.90, yielding the final training set of 3,260 structures (3260_training.zip). All structural alignments were performed using C4′ atoms and the default US-align settings for RNA.

To estimate the statistical potential, we aggregated interaction counts from the fixed training set of 3,260 RNA structures using template-centered alignment groups (Fig. 1a). These groups were used solely for statistical aggregation and did not change the composition of the training set. We performed all-against-all structural comparisons using US-align^60^, with each training structure serving in turn as a template. Each group included the template itself and all other training structures with a C4′-based TM-score of at least 0.45 to that template. This procedure defined 3,260 potentially overlapping alignment groups, as a structure could meet the inclusion criterion for multiple templates. Within each group, only residues aligned to the template contributed to the interaction-count tensors; unaligned regions were masked. Each structure was assigned a weight equal to the number of its aligned nucleotides divided by the longest aligned length in that group. For each tensor bin, the counts from individual structures were multiplied by their respective weights, summed across group members, and divided by the sum of the weights. Each group therefore contributed a single alignment-length-weighted average count tensor to the global statistics, rather than the unnormalized sum of its members’ counts.

### Sequence generation algorithm in DS3dRNA

#### Input construction

DS3dRNA takes RNA tertiary-structure information as its principal input (Fig. 1b,c). The design target comprises one or more 3D RNA scaffolds in PDB format. A single scaffold defines a single-state design task, whereas two or more residue-matched conformations define a multistate task in which the same sequence is optimized against all input structures. Multi-state targets can be obtained from experimentally determined ensembles such as solution NMR models, alternative functional conformations such as apo and holo riboswitch structures, or predicted structures generated by AlphaFold3^31^ or RoseTTAFoldNA^61^. DS3dRNA also supports mixed inputs comprising a full-length target scaffold and one or more shorter 3D motifs or fragments: each auxiliary fragment is mapped to user-specified residues of the full-length target; positions outside the mapped region are masked. This masking scheme allows unequal-length fragments to contribute local structural restraints without requiring artificial coordinates for unspecified regions. To preserve continuous and physically plausible geometry, motifs or fragments should first be superposed or assembled onto the target using US-align^60^ or RNA template-assembly methods such as Vfold^59^, FARFAR2^17^, 3dRNA^62^, RNAComposer^54^ or FebRNA^50^.

Secondary-structure constraints can be supplied to DS3dRNA in dot-bracket notation. For targets with available all-atom structures, base-pair assignments can be extracted using x3DNA-DSSR^56^ (v.1.9.10) before coarse-graining. In our benchmarks, only the resulting base-pair assignments were passed from this annotation step to the design procedures that used secondary-structure constraints; native nucleotide identities and nucleotide-specific all-atom annotations were not transferred. This preprocessing therefore provided target pairing constraints rather than reference sequences for design. In automatic mode, DS3dRNA infers base-pairing constraints, including pseudoknotted pairings, from inter-bead distances and base-anchor orientations in the target P, C4′ and N1/N9 scaffold. During sequence optimization, nucleotide substitutions at constrained paired positions are restricted to A–U, G–C or G–U combinations.

DS3dRNA also allows users to specify fixed nucleotide identities at selected sequence positions. These identities remain unchanged throughout sequence optimization, while the remaining positions are redesigned. The choice of fixed positions and nucleotide identities may be informed by experimental measurements, such as deep mutational scanning (DMS)^21^, evolutionary conservation in RNA multiple sequence alignments, or sequence preferences inferred from nucleic acid language models^30,63,64,65^ and RNA family-specific generative models^26,45,46,66^. These optional hard constraints enable experimental, evolutionary and model-derived priors to be incorporated into structure-guided sequence design.

#### Higher-order energy-based Markov chain Monte Carlo sampling

To generate sequence ensembles, we developed a Markov chain Monte Carlo^58^ workflow driven by the three-body interaction potential TriRNASP^19^ (Supplementary Fig. S1). After encoding the structural, secondary-structure and frozen-site constraints, DS3dRNA initializes an ensemble {***S*_0_**_}_ of 320 sequences by assigning A, U, C and G with equal probability at each unconstrained position. At Monte Carlo step t, every sequence *S_t,i_* _i_s scored with the TriRNASP-Rough potential. The resulting energies define a Boltzmann-weighted^67^ categorical distribution for the nucleotide at each position *I*:

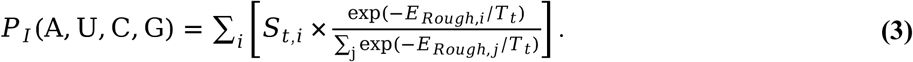

In Eq. (3), *S_t,i_* _i_s the one-hot nucleotide vector at position I in sequence i, *E_rough,i_* _i_s its Rough energy and T_ₜ_ is the annealing coefficient at step t. T_ₜ_ was decreased exponentially from 100 to 4.5 k_BT_ over the trajectory. At each step, 22 sequences were independently sampled from the position-specific distributions and combined with the ten lowest-Rough-energy members of the current 320-sequence ensemble, yielding a batch of 32 candidates. This batch size was chosen to enable efficient parallel evaluation on CUDA-enabled NVIDIA GPUs using PyTorch v.2.7.1+cu128. Detailed CPU–GPU runtime benchmarks are presented in Supplementary Fig. S7.

The 32 candidates were screened using a restricted Turner 2004 nearest-neighbor thermodynamic model for RNA^47,68,69,70^ and the SantaLucia nearest-neighbor framework for DNA^48,71,72,73^. We retained local energetic terms and excluded multibranch-loop, exterior-loop, coaxial-stacking and dangling-end contributions, which depend more strongly on global secondary-structure context^47,70,74^. The five candidates with the lowest minimum free energies under this restricted model were rescored with the TriRNASP-Fine potential. The lowest-Fine-energy candidate was then proposed for a Metropolis^58^ update:

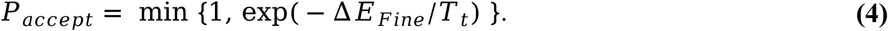

In Eq. (4), ΔE_fine i_s the Fine-energy difference between the proposed and current sequences. If the proposal was accepted, it was added to the sequence collector and used as the parent for batch mutation to generate the next 320-sequence ensemble; if it was rejected, the current sequence was retained as the parent. Every ten Monte Carlo steps, the per-position mutation probability P_mut w_as adjusted using the acceptance fraction over the preceding ten steps: P_mut w_as increased when the fraction exceeded 0.5, decreased when it was below 0.1 and otherwise left unchanged.

After a preset number of Monte Carlo steps (10,000 by default), the sequences accepted during the final 40% of the trajectory were pooled to estimate a position-specific nucleotide distribution, *Q_I_*_(_A, U, C, G), for each position I. An output decoder then applied dynamic programming to recover the sequence maximizing the cumulative log-probability under *Q_I_*, subject to the constraint that no homopolymer run exceed five consecutive nucleotides. This constraint limits long homopolymeric tracts and serves as a lightweight negative-design heuristic against repetitive sequences. More generally, experimental RNA design studies support combining energy-based objectives with sequence-level design rules and iterative functional validation^12,27,75^. Penalties on repeated n-mers were similarly proposed as a complement to standard nearest-neighbor energetic models in a large-scale Eterna study^75^, and the same maximum-run constraint is imposed in the Eterna OpenKnot challenge (https://eternagame.org/challenges/11843006). DS3dRNA additionally records sampled sequences and their energies during each design run. In batch-design mode, candidate sequences are ranked by their TriRNASP-Fine energy using DS3dRank, the ranking module of DS3dRNA. DS3dRank further supports high-throughput (approximately one million sequences per second on an NVIDIA RTX 5080 GPU) fixed-backbone ranking of arbitrary externally supplied sequences—for example, candidates designed by gRNAde^14^ or sequences with measured DMS-based fitness^21^.

### Benchmark curation and redundancy control

#### Construction of the non-redundant T83 benchmark

To establish a common benchmark for all methods, we evaluated each method from its public release without retraining, using the default settings provided by its authors. We collected the available training, validation and test sets for RhoDesign^13^, gRNAde^14^, R3Design^41^, RiboDiffusion^42^, RIdiffusion^43^, AlignIF^44^ and DS3dRNA. Before duplicate removal, the combined training pool contained 27,014 native coordinate files representing 6,414 unique PDB identifiers (Train_pdb_id_all_methods_collection_6414.txt). RhoFold-generated decoys used to train RhoDesign and RiboDiffusion were not included because the corresponding coordinates were not fully publicly available. The combined validation and test sets represented 999 unique PDB identifiers (Test_pdb_id_all_methods_collection_999.txt) and constituted the starting pool for benchmark construction.

For each of the 999 PDB identifiers, we downloaded the complete PDBx/mmCIF archive from the RCSB PDB^76^ (https://www.rcsb.org/downloads) and converted the coordinates to PDB format using Gemmi v.0.6.4^77^. Multi-model entries were separated into individual coordinate models, yielding 1,575 files. To avoid disrupting native interchain contacts, we annotated interchain base pairs with x3DNA-DSSR^56^ and separated chains only when no interchain base pairing was detected, and treated each retained group of mutually paired chains as one structural unit. This procedure produced 1,905 structural units, of which 1,584 were within the predefined length range of 16–512 nucleotides.

We next performed all-against-all structural comparisons using US-align^60^ with C4′ atoms and the default RNA settings. A pair was considered redundant only when both the TM-score and the sequence identity over the structurally aligned region were ≥0.90, and one representative was retained from each redundant group. This structural and aligned-sequence filter reduced the candidate pool to 1,309 molecules. Candidate sequences were then compared with the training-set sequences derived from all 27,014 coordinate files using CD-HIT^78^ v.4.8.1, with sequence-identity and alignment-coverage thresholds of 0.90. Excluding all matches yielded 83 molecules, which constituted the non-redundant T83 benchmark.

As an independent sequence-level check, we used Parasail^79^ v.1.3.4 with alignment-coverage thresholds of 0.90 to calculate the maximum sequence identity between each T83 molecule and the combined training pool. For multichain molecules, chain-level identities were weighted by chain length. The resulting maximum identities ranged from 0.60 to 0.90 (T83_molecule_report_vs_train.xlsx).

#### RNA-only inverse-folding benchmark T25

Sequence recovery on a fixed scaffold measures agreement with native nucleotide assignments but does not establish whether a designed sequence folds to the target 3D structure. We therefore constructed an RNA-only inverse-folding self-consistency benchmark to assess target-fold recovery while reducing confounding interactions with protein or ligand partners. From T83, we retained seven RNA-only structural units: chains A and B of PDB 1BR3; chains A–D of 1EGK; and chains A and B of 1FIX, 1JZV, 3SSF, 5CKK and 7PDU.

To broaden the structural diversity of the benchmark, we additionally included 18 publicly released structures from the CASP15 and CASP16 target lists whose target type was annotated as RNA or NucA: PDB 8BTZ, 7PTK, 7PTL, 8UYS, 8UYE, 8UYG, 8UYJ, 9CFN, 9C2K, 9ELY, 9MEE, 9CBU, 9CBX, 9ZC7, 9ZC8, 9MCW, 9J6Y and 9C75. We excluded any target whose PDB identifier occurred among the 6,414 unique identifiers in the combined training pool of the benchmarked methods. The resulting 25-target dataset, comprising seven targets from T83 and 18 targets from CASP15 and CASP16, was designated T25 (T25.zip).

#### The multistate design benchmark T11

We constructed T11 to compare DS3dRNA and gRNAde^14^, the two methods evaluated in this study that support sequence design jointly conditioned on multiple backbone conformations. T11 comprised 11 target ensembles containing 168 coordinate models. Each ensemble was treated as a single design target, with one sequence designed jointly against all constituent conformers; individual conformers were not treated as independent targets.

We searched the RCSB Protein Data Bank^76^ (PDB; accessed 1 December 2025) using the keyword “riboswitch” and the experimental-method filter “SOLUTION NMR”. The search yielded ten entries (PDB IDs 2KXM, 2KZL, 2L1V, 2LI4, 2MIY, 2MXS, 2N0J, 5KH8, 6HAG and 9HRO), representing RNAs of 27–59 nucleotides. All deposited conformers were retained, yielding ten NMR ensembles with 156 coordinate models. To include a longer RNA target with complete residue-level coordinate coverage, we added 12 crystal structures of the 174-nucleotide sensing domain of the Thermotoga maritima lysine riboswitch reported by Serganov et al.^7^ (PDB IDs 3DIG, 3DIL, 3DIM, 3DIO, 3DIQ, 3DIR, 3DIS, 3DIX, 3DIY, 3DIZ, 3DJ0 and 3DJ2). These structures were grouped into a single target ensemble. None of the selected PDB entries was included in the training set of either DS3dRNA or gRNAde. T11 is available as T11.zip.

#### CASP17 P20 kissing-multiloop benchmark

We considered four computationally designed CASP17 RNA targets with PDB structures available at the time of analysis: R2304 (11EH), R2305 (11AG), R2306 (10ZU) and R2307 (10ZT). Three entries—10ZT, 10ZU and 11EH—were monomeric cryo-electron microscopy structures of independent sequences designed for the P20 “Kissing Multiloops” target in Round 3 of the Eterna OpenKnot challenge^12^. The remaining entry, 11AG, was an MPNN–RFdiff-designed dimer and was excluded from the monomeric comparison.

Pairwise US-align comparisons using C4′ atoms yielded TM-scores of 0.432 for 10ZU–10ZT, 0.504 for 10ZU–11EH and 0.522 for 10ZT–11EH, indicating differences among the experimentally resolved folds despite their shared secondary-structure design target. We selected 10ZU, the 4.0-Å cryo-EM structure of an RNA designed by the fixed-backbone message-passing model MPNN-fixbb, as the reference scaffold. This provided a focused test of DS3dRNA on an experimentally resolved fold originating from neural inverse design. MPNN-fixbb had also shown experimental design capability in an earlier round of the OpenKnot challenge, with Townley et al. reporting improved average performance over the W03 starting sequence in Round 1^12^.

All DS3dRNA designs for 10ZU were generated in DS3dRNA_auto mode using only the sequence-masked coarse-grained scaffold represented by P, C4′ and N1/N9 atoms. Neither the original nucleotide identities nor the OpenKnot target secondary structure, including its seven prescribed stems, was supplied; base-pairing constraints were inferred automatically from the scaffold geometry. For each design trajectory, we retained both the decoded per-run output sequence and the sequence with the lowest Fine energy, denoted DS3dRNA_auto and DS3dRNA_auto_Emin, respectively. All other benchmarked methods were run using their publicly released implementations with default settings, following the authors’ design instructions.

### Evaluation metrics and benchmark protocol

#### Benchmark execution and input conditions

All comparison methods were evaluated using the authors’ released checkpoints and inference implementations, without retraining or benchmark-specific hyperparameter tuning. The documented inference procedures were retained, with the sequence count and input representations configured as described below. The evaluated architectures, secondary-structure inputs and method-specific processing steps are summarized in Table 1. Three DS3dRNA configurations were evaluated: DS3dRNA, which used a 3D scaffold without secondary-structure constraints; DS3dRNA_auto, which additionally applied base-pairing constraints inferred by the internal coarse-grained geometric parser; and DS3dRNA_SS, which used externally supplied secondary-structure constraints. DS3dRNA and DS3dRNA_auto therefore required the same external scaffold input but differed in whether internally inferred pairing constraints were used during sequence optimization.

For methods accepting external secondary structure, a common base-pair annotation was extracted from the target structure using x3DNA-DSSR v.1.9.10-linux^56^. Only the resulting pairing assignments were transferred through this annotation interface, with conversion to the representation required by each method, such as dot-bracket notation or a contact map. Native nucleotide identities and nucleotide-specific all-atom annotations were not transferred through this step. No external DSSR annotation was supplied to DS3dRNA_auto or R3Design, which used their internal geometric and ModeRNA-based^80^ procedures, respectively. For gRNAde^14^, the target secondary structure was supplied as required by the released design workflow used in this study. AlignIF^44^ retained its native MStA retrieval procedure. RIdiffusion^43^ used the authors’ PDB-only inference pathway without an auxiliary.st file containing per-residue secondary-structure-element labels. For R3Design^41^, multichain scaffolds were converted to a single-chain representation before inference so that all target residues, rather than only the first labeled chain, were processed together. The original coordinates and target residue order were retained.

#### Sequence generation and benchmark targets

Design calculations were performed on a workstation equipped with an NVIDIA GeForce RTX 5080 GPU. Each evaluated method or configuration generated 100 sequences per benchmark target. In the single-state benchmarks, one target corresponded to one reference RNA scaffold. The T11 multistate comparison was restricted to DS3dRNA and gRNAde. For each T11 target, 100 sequences were designed jointly against the complete conformational ensemble; the number of input conformers matched the actual number in that ensemble. Individual conformers were not treated as separate sequence-design targets.

#### Consensus candidates and energy-selected designs

For structure-based self-consistency evaluation, one consensus candidate was constructed per method and target from the generated sequence collection. For DS3dRNA, the target-level consensus used the same underlying nucleotide-frequency rule and homopolymer cap as the per-run output. The target-level profile was decoded from left to right by selecting the highest-frequency admissible nucleotide, with a maximum of five consecutive identical nucleotides. For the benchmark inputs, the homopolymer count was reset at chain boundaries. In the released target-level processing script, duplicate output sequences were weighted by their count, and records with Fine energy more than 10,000 k_BT_ above the minimum for that target and configuration were excluded before consensus construction. Consensus candidates for comparator methods were obtained from their generated sequences using the position-wise frequency procedure. A consensus candidate was not required to coincide with an individual generated sequence. The public post-processing utility the Script and usage examples are provided in the repository’s Examples directory.

For DS3dRNA, the suffix _Emin denotes the lowest-Fine-energy sequence selected from the target-level output collection for the corresponding input configuration. Sequence recovery and MacroF1 for an _Emin entry were calculated for this single selected sequence, and structural predictions evaluated the same candidate. Ordinary DS3dRNA sequence summaries instead used the retained per-run outputs, as specified below. Consensus and _Emin structural results were kept separate; they were not combined by selecting the better-performing candidate for each target.

#### AlphaFold3 prediction protocol

Quantitative structural analyses were performed using the official local AlphaFold3 (v.3.0.1)^31^ implementation on the Wuhan University high-performance computing platform equipped with NVIDIA A100 GPUs. To assess cross-platform reproducibility and facilitate replication through a publicly accessible workflow, selected reported candidates were also submitted to AlphaFold Server^31^. The same RNA sequences, chain composition and copy numbers were used for corresponding local and server runs, for both single-chain and multichain targets. No target coordinates, target secondary structure or user-specified structural restraints were supplied.

Local inference used the default settings of ten recycling iterations and five diffusion samples, consistent with the server configuration used in this study. A single random seed was fixed at 1 for both platforms, and the standard data-processing pipeline was retained. All five local predictions were evaluated separately, and C4′ RMSD and C4′ TM-score values were averaged over the five predictions for each candidate–reference comparison, without best-of-five selection.

Server reruns reproduced the corresponding local structural results, providing a check that the reported predictions could be reproduced through the public server without access to our HPC environment. Only local predictions were included in the quantitative benchmark; server outputs were used solely for the reproducibility check and were not pooled with the local results. The structural prediction results are available in AlphaFold3_results_All.zip. These analyses assessed predicted structural self-consistency rather than experimentally validating the designed folds.

#### Sequence-level metrics

For each target t, let ℐ_t_ denote the set of evaluated residue positions and N_t_ its size. The reference sequence and the j-th designed sequence are denoted by 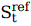 and 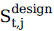, respectively. Sequence recovery was defined as

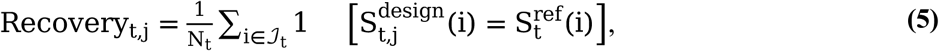

where the indicator equals 1 for a nucleotide match and 0 otherwise. Each evaluated position contributed once, including positions in all target chains. For a multistate target, each design was compared once with the shared reference sequence, irrespective of the number of conformers.

MacroF1 was calculated across the nucleotide classes represented in the evaluated reference sequence:

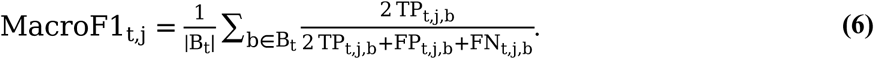

Here, B_t_ is the subset of {A, U, C, G} present at the evaluated reference positions, and TP, FP and FN are the position-wise true-positive, false-positive and false-negative counts for nucleotide class b. Classes absent from the reference were excluded. A represented class that was never assigned in a design received an F1 score of zero. This target-specific definition was used consistently for all methods.

#### Structure-level metrics

**C4′ RMSD**. Native and predicted C4′ atoms were matched by target chain and residue order. One optimal rigid-body transformation was fitted to the complete set of matched coordinates using Bio.PDB.Superimposer in Biopython v.1.85^81^. RMSD was reported in ångströms. For multichain targets, the fit was performed over the whole molecule rather than separately for each chain.

**C4′ TM-score**. Global fold similarity was calculated using US-align v.20241108^60^, with explicit C4′-atom selection and the RNA-specific normalization scale:

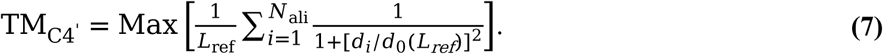

L_ref_ is the native-target length used for normalization, N_ali_ is the number of structurally aligned residue pairs, d_i_ is the C4′ distance for aligned pair i under the US-align superposition, and d_0(_L_ref_) is the RNA-specific length-dependent scale. The native-target-normalized score was used throughout. For US-align scoring of multichain molecules, all target residues were arranged in a consistent target order and represented as a single continuous chain in both the native and predicted coordinate files. This allowed the entire molecule to be aligned in one comparison without retaining separate chain labels in the scoring input. Coordinates were unchanged by this relabelling.

RMSD used the prescribed target residue correspondence, whereas TM-score used the structural alignment returned by US-align.

#### Target-level and dataset-level aggregation

For ordinary sequence-design outputs, sequence recovery and MacroF1 were first summarized within each target and then averaged across targets without weighting by sequence length or conformer count. DS3dRNA target-level means used the retained per-run outputs after the Fine-energy filter described above, with duplicate sequences weighted by count, i.e., this is equivalent to averaging the retained run outputs. In contrast, each _Emin entry contributed the score of one minimum-energy sequence per target. Comparator target means were calculated across their 100 generated sequences. Every target contributed equally to the dataset-level metrics, and for T11, the averaging units were its 11 ensembles, not the individual conformers.

For single-state structural benchmarks, the five AlphaFold3 predictions were first averaged at the candidate–target level, followed by an unweighted mean across targets. Consensus and _Emin candidates were aggregated separately. The one-to-many comparisons for the long T11 target remained an illustrative case analysis and were not included in a T11-wide structural average.

## Experimental methods

### De novo design of the Mango II G-quadruplex aptamer

RNA G-quadruplexes (G4s) comprise stacked guanine quartets stabilized by Hoogsteen hydrogen bonding and monovalent cations^5,82,83^, particularly K^+^. Conventional base-pair-based secondary-structure annotations do not explicitly describe the Hoogsteen interactions, quartet organization or stacking geometry of the G4 core. We selected the fluorogenic aptamer Mango II (PDB ID: 8VY0)^5,9,53^ as a G4 design target to evaluate sequence design using 3D structural information beyond conventional secondary-structure constraints.

DS3dRNA was supplied with the experimentally determined RNA 3D scaffold and its corresponding secondary structure. We performed 100 independent design trajectories, ranked the resulting sequences by design energy and selected the three lowest-energy sequences (DS-1, DS-2 and DS-3) for experimental evaluation. RhoDesign^13^ was run in its two modes: 3D scaffold plus secondary structure (Rh-1) and 3D scaffold only (Rh-2). One sequence was generated in each mode at a sampling temperature of 1 × 10^−5^. All five designed sequences were evaluated alongside wild-type (WT) Mango II by fluorescence titration^5,53^.

#### RNA annealing and fluorescence titration

Unlabelled RNA at 5 µM was annealed in 10 mM Tris (pH 7.0), 140 mM KCl and 1 mM MgCl_2 b_y heating at 95 °C for 10 min, cooling on ice for 15 min and equilibrating at room temperature for 30 min.

Fluorescence titrations were performed using a Cytation 3 Imaging Reader in 10 mM Tris (pH 7.0), 140 mM KCl, 1 mM MgCl_2 a_nd 0.005% Tween-20. The final total RNA concentrations were 0, 0.05, 0.1, 0.2, 0.5, 1, 2, 5, 10, 20, 50 and 100 nM. TO1-3PEG-Biotin was added after RNA dilution to a final concentration of 1 nM in a total assay volume of 200 µL. Fluorescence was recorded at excitation and emission wavelengths of 510 and 535 nm, respectively. Three replicate titration series were measured for each RNA.

#### Binding analysis

Binding was analyzed using the tight-binding quadratic equation^9,53,84^:

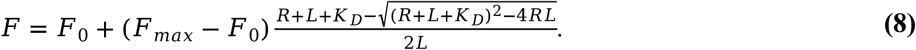

Here, F is the measured fluorescence; R and L are the total RNA and TO1-3PEG-Biotin concentrations, respectively, with L held at 1 nM; F_0 i_s the fluorescence measured at zero RNA concentration in the corresponding replicate; F_max i_s the fitted fluorescence at saturation; and K_D i_s the apparent dissociation constant. Each replicate was fitted separately by unweighted nonlinear least squares, with F_0 f_ixed to its measured value. The fitted parameters were F_max a_nd the base-10 logarithm of K_D_, ensuring a positive K_D._ Apparent K_D v_alues were reported as the mean ± s.d. of the three replicate estimates (n = 3).

#### Comparison of fluorogenic responses

To compare RNA-induced fluorescence activation among the designed aptamers and WT Mango II under matched assay conditions, fluorescence at each RNA concentration was normalized to the measured zero-RNA, dye-only baseline from the same replicate by dividing by this baseline and subtracting one. Normalized responses were summarized as the mean ± s.d. of three replicates. The mean normalized titration curve was fitted to the normalized form of Eq. (8), with a fixed zero intercept and a fitted saturation amplitude. Weighted nonlinear least squares was performed using pointwise replicate s.d. values as absolute uncertainties. The zero-RNA point was excluded from the weighted residuals because its normalized response and s.d. were zero by construction.

The fitted amplitude, reported as “Saturation Normalized RFU”, quantified the baseline-relative fluorescence increase at saturation and was used as a comparative measure of fluorogenic response. Adding one to saturation amplitude yielded the corresponding assay-specific fluorescence-enhancement ratio. Standard errors for saturation amplitude were obtained from the fitted covariance matrix, treating the measured normalization baselines as fixed. Saturation amplitudes were estimated from the mean-curve fits, whereas apparent K_D v_alues were obtained from the separate replicate fits described above, providing complementary assessments of fluorescence activation and binding affinity. Apparent K_D v_alues and saturation amplitudes were reported as not determined (n.d.) when no well-resolved saturating response was observed within the tested concentration range.

### De novo design of ES2 twister ribozymes from a predicted structural ensemble

Catalytic RNAs provide a stringent test of structure-based design because their activity depends on precisely arranged 3D active sites^20^. The twister fold contains a compact double-pseudoknot core in which conserved nucleotides form canonical and noncanonical contacts that position the scissile phosphate and catalytic nucleobases^8,20,85^. Cleavage proceeds through an in-line transesterification reaction involving activation of the 2′-hydroxyl nucleophile and general acid–base catalysis; divalent ions also contribute to folding, conformational organization and catalytic competence^20,85^. Cleavage activity therefore provides a functional readout of the ability of a designed sequence to assemble a catalytically competent tertiary structure.

Experimentally resolved RNA structures represent only a small fraction of the sequence space catalogued in resources such as RNAcentral^23^ and Rfam^86^. Structure-based design must therefore also accommodate predicted scaffolds, particularly when the exact construct used in a functional assay lacks an experimental structure. We selected the ES2 twister system, a bimolecular ribozyme derived from the P3-type env-9 environmental sequence and comprising separate ribozyme and substrate strands^8,52^. An experimentally determined 3D structure matching the complete ES2 ribozyme–substrate construct used in the cleavage assay was not available in the Protein Data Bank. We therefore generated a predicted structural ensemble of the complete complex before sequence design.

#### Structural inputs and candidate selection

The ES2 complex was modeled using the AlphaFold Server^31^, with the wild-type ribozyme strand assigned to chain A and the substrate strand to chain B. Five Na^+^ ions and two Mg^2+^ ions were included in the input. Prediction used one model seed and five diffusion samples, yielding five models of the ribozyme–substrate complex. All five models were used jointly as structural targets for multistate DS3dRNA design. Secondary-structure annotations were extracted from the predicted structures using DSSR and supplied as additional design constraints.

Using this five-model ensemble and the associated secondary-structure constraints, we performed 100 independent DS3dRNA design trajectories. The resulting sequences were ranked using the DS3dRNA energy function, and the five lowest-energy sequences, designated DS-1–DS-5, were selected for experimental validation. Only the ribozyme strand was redesigned; the substrate sequence was held fixed throughout optimization and was identical to that used with the wild-type ES2 ribozyme. All designed ribozymes were therefore evaluated against the same substrate as the wild type, avoiding substrate-sequence changes as a confounding variable.

#### Ribozyme cleavage activity assay

Cleavage activities of wild-type ES2 and the designed ribozymes were measured using a procedure adapted from the previously reported ES2 twister assay^8^.

Wild-type or designed ribozyme RNA was mixed with 5′-Cy3-labeled substrate RNA (5′-Cy3-ACCAGCGGCAGAAUGCAGCUUUAUUGCC-3′) in annealing buffer containing 50 mM NaCl and 0.1 mM EDTA (pH 7.0). The ribozyme and substrate concentrations in the annealing mixture were 1 μM and 50 nM, respectively. Mixtures were heated at 80 °C for 5 min, cooled to 25 °C at approximately 0.5 °C min^−1^ and equilibrated at 25 °C for 20 min. Cleavage was initiated by adding an equal volume of 2× cleavage buffer, corresponding to calculated final ribozyme and substrate concentrations of 0.5 μM and 25 nM, respectively. The final reaction mixture contained 25 mM HEPES (pH 7.0), 10 mM MgCl_2_, 100 mM NaCl and 0.05 mM EDTA. Reactions were incubated at 25 °C for 20 s, after which a 2-μL aliquot was immediately transferred into 13 μL of stop solution containing 95% (v/v) formamide, 50 mM EDTA and RNA loading dye.

Cleaved and uncleaved substrate RNAs were resolved by 20% denaturing polyacrylamide gel electrophoresis (PAGE). Cy3 fluorescence was detected using a Typhoon fluorescence imaging system, and band intensities were quantified using ImageJ. The percentage of substrate cleaved at the 20-s endpoint was calculated as

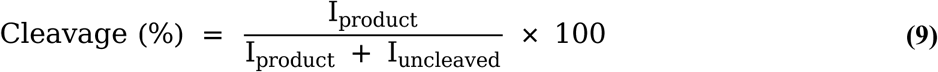

where *I*_product a_nd *I*_uncleaved a_re the fluorescence intensities of the cleaved product and uncleaved substrate bands, respectively. Each ribozyme was tested in four replicate experiments (n = 4).

#### RNA oligonucleotides

All RNA oligonucleotides for Mango II G-quadruplex aptamer, ES2 twister ribozymes and the substrate were synthesized by Sangon Biotech (Shanghai, China).

## Data availability

The structural datasets, benchmark collections and supporting prediction outputs are available in the DS3dRNA public data archive on Google Drive. The archive includes the training set, T83, T25, T11 and CASP17 P20 structures, the training and test PDB lists, the T83 similarity report and AlphaFold3 results. The complete target-level design-run profiles are provided as Supplementary_T25_T83_100run_profiles.pdf. Raw Mango II titration measurements and annotated twister source gels are provided in Supplementary Table S1 and Supplementary Fig. S8, respectively.

## Code availability

DS3dRNA source code and inference instructions are available at the DS3dRNA GitHub repository. The Examples directory contains single-state design, multistate design and sequence-ranking workflows. Repositories for the comparator implementations are listed in Table 1. Software and computational resources are listed by version, role and availability in Supplementary Table S4. Versioned benchmark tools and prediction settings are additionally specified in the relevant Methods subsections.

## Supporting information

DS3dRNA_SI

## Acknowledgements

We thank Professors Shi-Jie Chen (University of Missouri) and Jian Zhang (Nanjing University) for helpful discussions. This work was supported by the National Natural Science Foundation of China (grant nos. 12375038 and 12075171 to Z.-J.T.; 12374216 and 12074294 to X.-H.Z.; 12205223 to Y.-L.T.; and 12404246 to X.-C.Z.), Hubei Provincial Natural Science Foundation of China (No. 2024AFE008 and 2025AFA021 to X.-H.Z.). Parts of the numerical calculations were performed at the Supercomputing Center of Wuhan University.

## Author contributions

T.Y. conceived the study, developed the many-body interaction model, led the development of the DS3dRNA algorithm, implemented and tested the software, analyzed the computational and experimental results, and drafted the manuscript; D.W. performed the G-quadruplex aptamer experiments and contributed to their analysis and interpretation; X.-C.Z. performed the ribozyme experiments and contributed to their analysis and interpretation; X.-L.C. contributed to the secondary-structure thermodynamic modeling module, software implementation, testing, data analysis and manuscript discussion; H.-L.T., C.-C.Z. contributed to software testing, data analysis and manuscript discussion; Z.-J.T., X.-H.Z. and Y.-L.T. supervised the study, contributed to its conception and design, and participated in manuscript writing.

## Figure legends

**Extended Data Fig. 1.**
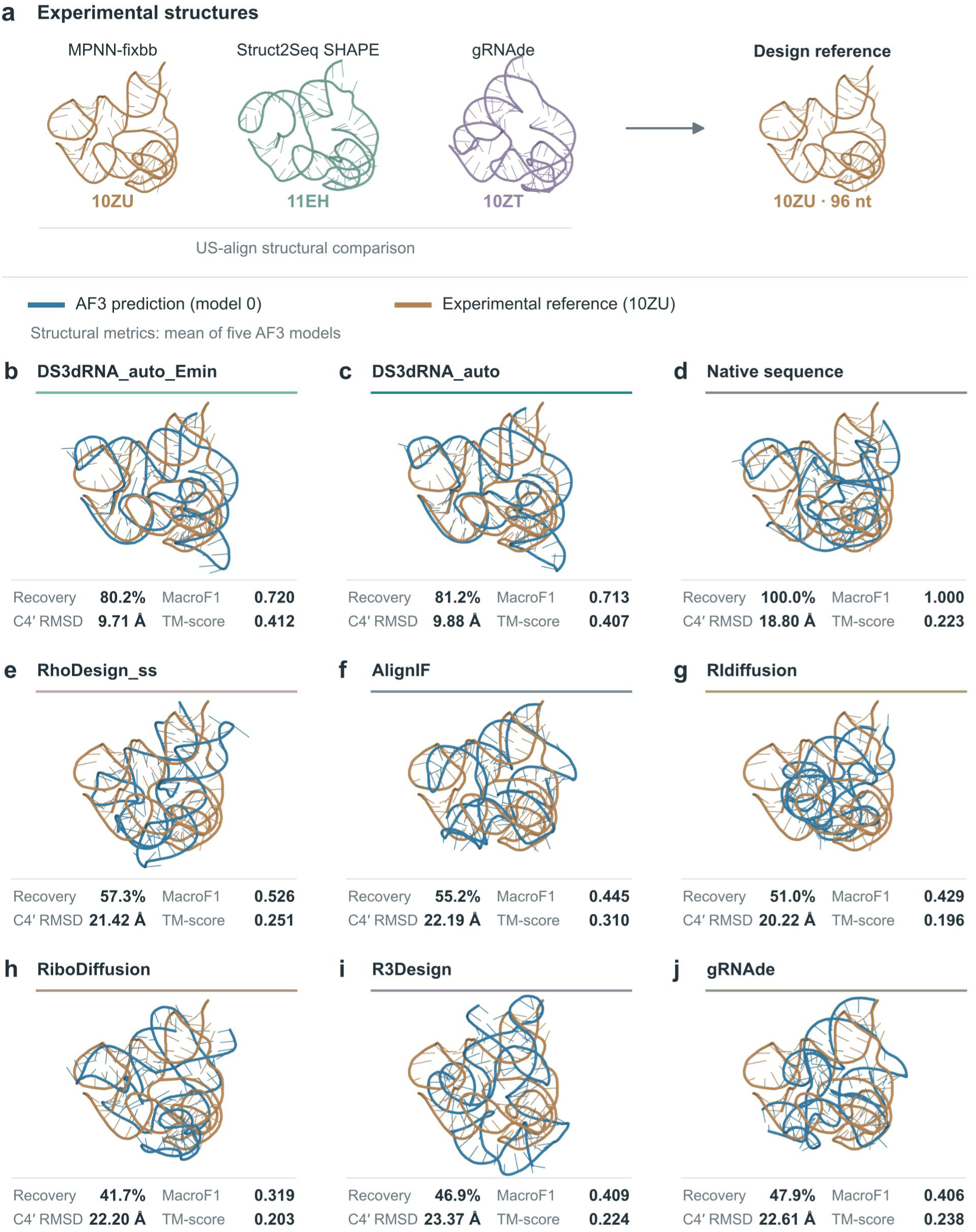
Sequence design for the CASP17 P20 kissing-multiloop scaffold. **a**, Experimental RNA structures associated with MPNN-fixbb (PDB 10ZU), Struct2Seq SHAPE (11EH) and gRNAde (10ZT), shown separately after US-align structural alignment. The modeled 96-nt region of 10ZU was used as the design reference. b–j, AlphaFold3 predictions for DS3dRNA_auto_Emin (b), DS3dRNA_auto (c), the native sequence (d), RhoDesign_ss (e), AlignIF (f), RIdiffusion (g), RiboDiffusion (h), R3Design (i) and gRNAde (j). Each panel shows model 0 in blue superposed on the experimental 10ZU reference in brown, with a common reference orientation and scale. Model 0 is used as a fixed-index illustration, without selection for structural accuracy. Reported C4′ RMSD and TM-score values are arithmetic means over all five models (indices 0–4), rather than values for the displayed model alone. C4′ RMSD uses least-squares fitting of all 96 residue-order-matched C4′ atoms, whereas TM-score uses the US-align structural alignment. Sequence recovery and MacroF1 were recalculated from the sequence encoded in the prediction files relative to the 96-nt reference. All five models within a method encode the same sequence; sequence metrics therefore describe one candidate rather than five independent designs.

