## Supplementary material for "De novo design of functional RNAs through higher-order interactions": DS3dRNA_SI

### through higher-order interactions

**This document accompanies the study and contains**

Supplementary Figs. S1–S8,

Supplementary Tables S1–S4,

Supplementary Data 1–3,

A note on retained design-run records and Supplementary References.

#### Supplementary Fig. S1

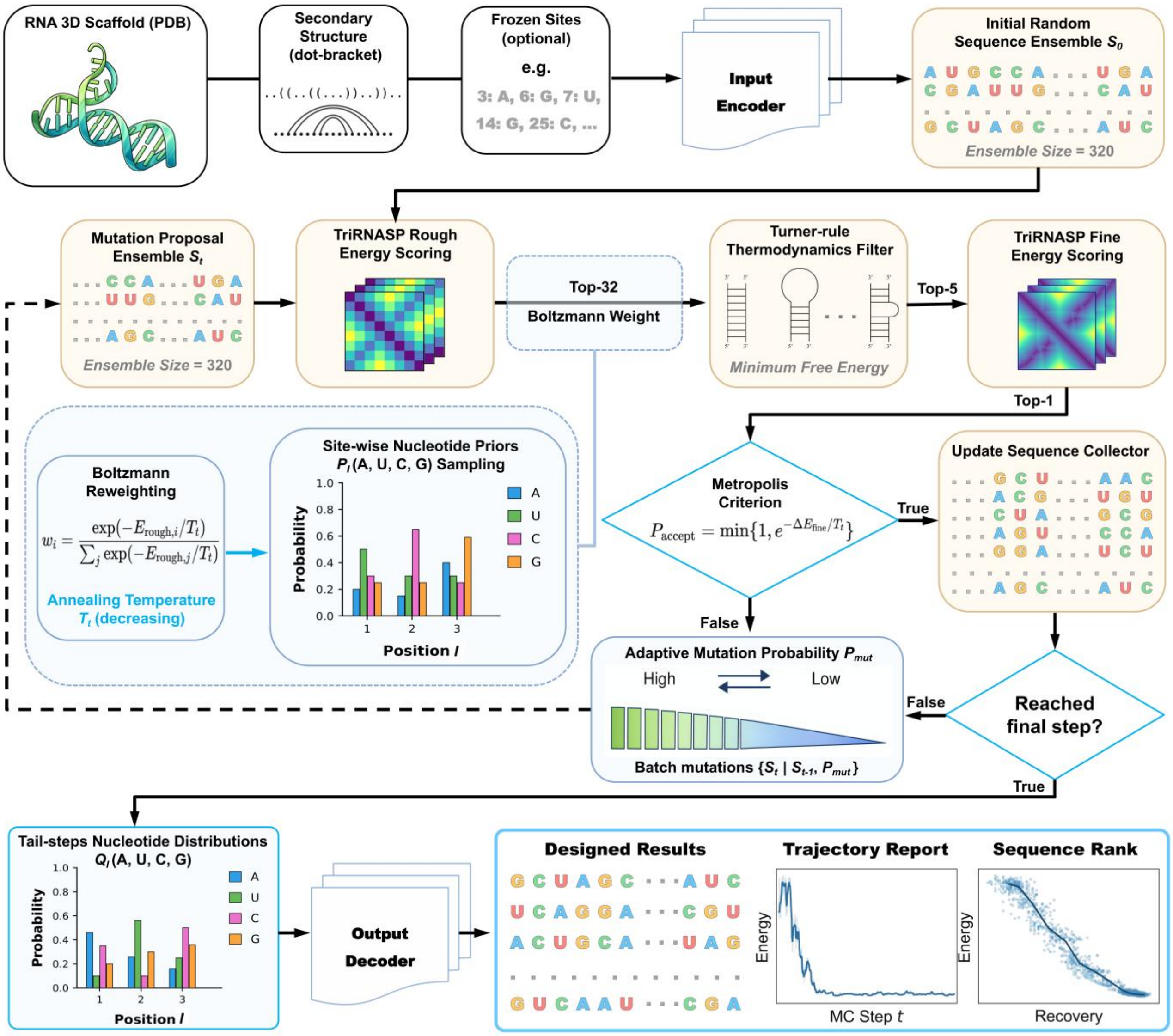


**Supplementary Fig. S1 | Sequence optimization by hierarchical energy screening and adaptive Monte Carlo sampling.** DS3dRNA accepts one or more residue-matched three-dimensional RNA scaffolds, optional secondary-structure constraints and optional fixed nucleotide identities. An initial ensemble of 320 sequences is scored with the coarse TriRNASP-Rough potential[1]. Boltzmann reweighting[2] defines position-specific nucleotide probabilities, which are used together with low-energy members of the current ensemble to form 32 candidate sequences. A restricted nearest-neighbor thermodynamic screen[3,4] retains five candidates for rescoring with TriRNASP-Fine. The lowest-Fine-energy candidate is proposed for a Metropolis update[5], and the accepted sequence, or the retained current sequence following rejection, seeds the next mutation ensemble. The annealing coefficient decreases during sampling, while the mutation probability is adjusted according to recent acceptance rates. After the final step, nucleotide frequencies from the retained tail of the run are decoded to produce an output sequence; low-energy candidates and recorded run profiles are also retained. The displayed nucleotide distributions and output plots are schematic. Algorithmic parameters and constraint handling are described in Methods.

#### Supplementary Fig. S2

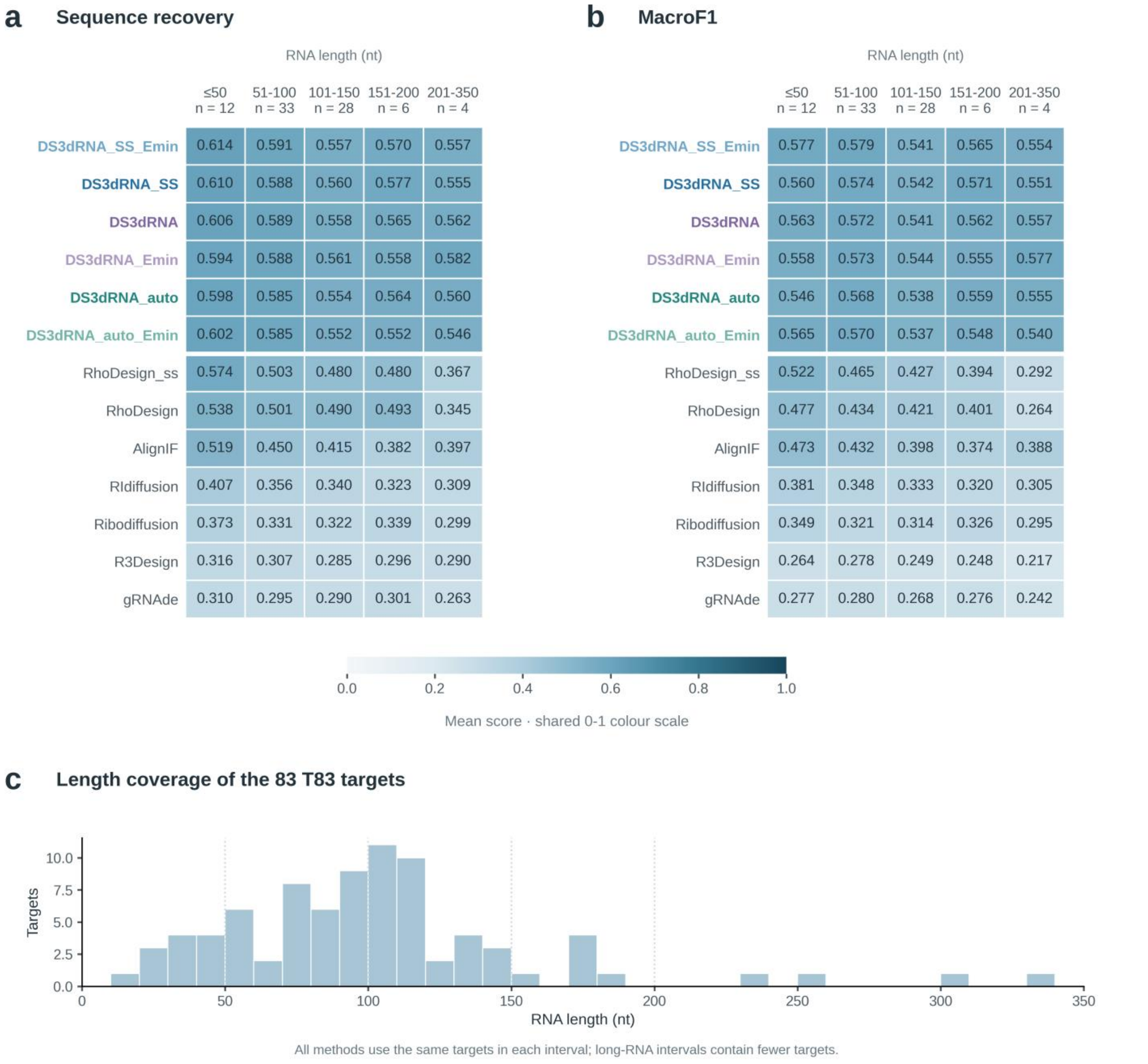


**Supplementary Fig. S2 | Sequence-design performance across RNA lengths in T83.** a,b, Mean native-sequence recovery (a) and macro-averaged F1 score (MacroF1; b) for the three DS3dRNA input configurations and their _Emin outputs and seven comparator configurations[6,7,8,9,10,11] shown in Fig. 2. Rows follow the Fig. 2 method order, and both heatmaps use the same 0–1 color scale. Values are arithmetic means across targets within each length interval; each ordinary DS3dRNA entry summarizes retained runs within a target, whereas each _Emin entry scores one minimum-energy candidate per target. The intervals ≤50, 51–100, 101–150, 151–200 and 201–350 nucleotides contain 12, 33, 28, 6 and 4 targets, respectively; all methods use the same targets in each interval. c, RNA-length distribution of the 83 T83 targets. Vertical dotted lines mark the length-interval boundaries used in a and b. The small numbers of targets in the two longest intervals should be considered when interpreting their means.

### Supplementary Fig. S3

#### Individual methods across RNA lengths

Every point is a target; lines join within-bin means at their observed mean lengths.

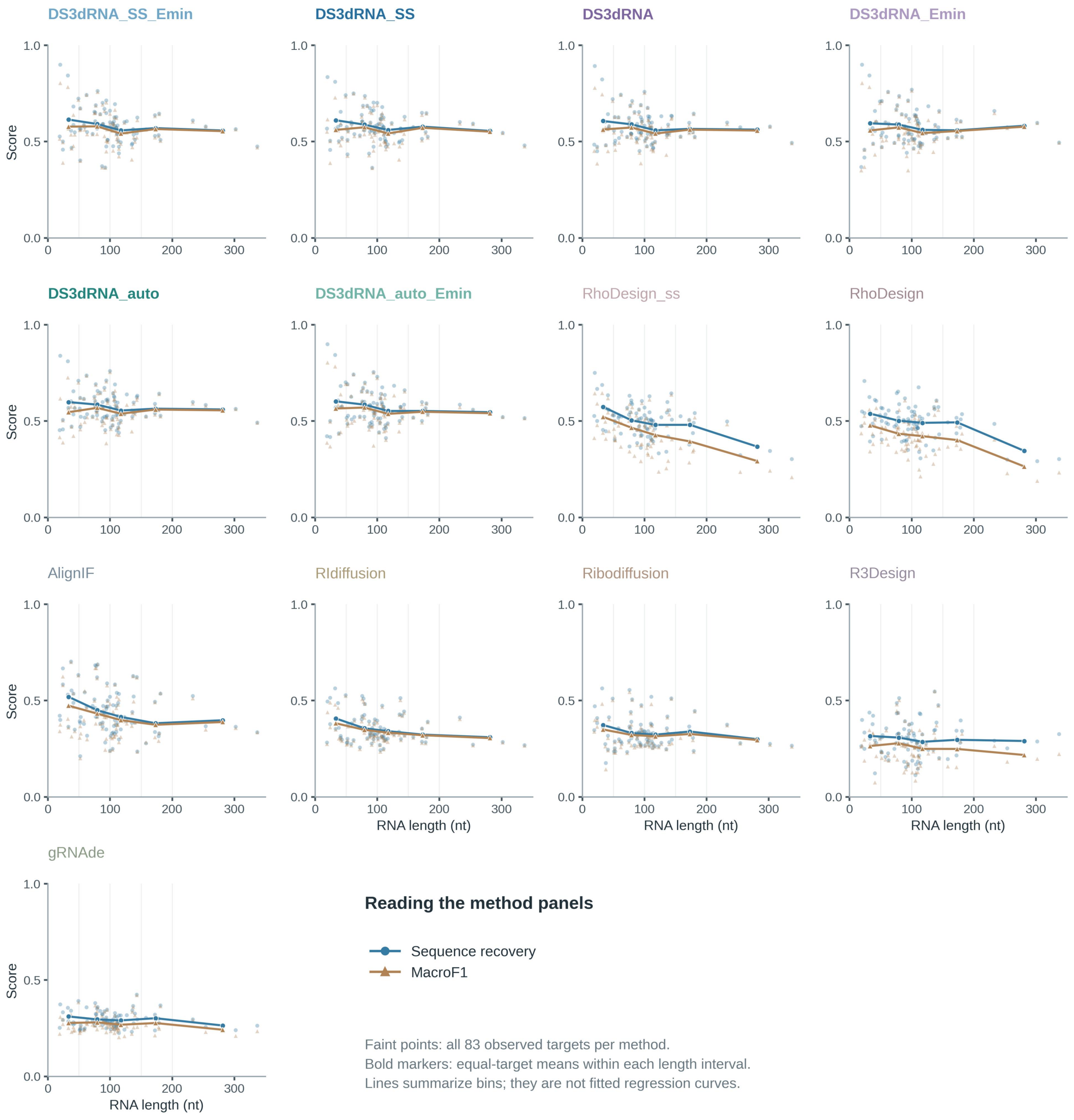


Length intervals: ≤50 / 51-100 / 101-150 / 151-200 / 201-350 nt; n = 12 / 33 / 28 / 6 / 4.

**Supplementary Fig. S3 | Target-level sequence recovery and MacroF1 as a function of RNA length.** Each subplot shows one method or input configuration on the same 83 T83 targets, in the order used in Fig. 2 and Supplementary Fig. S2. Small blue circles show sequence recovery and small ochre triangles show MacroF1 for individual targets, using retained-run means for ordinary DS3dRNA outputs and single-candidate scores for _Emin outputs. Larger markers indicate the arithmetic mean within each of the five length intervals, positioned at the observed mean RNA length of that interval. Lines connect these descriptive means; they are not fitted regressions or extrapolations. The intervals ≤50, 51–100, 101–150, 151–200 and 201–350 nucleotides contain 12, 33, 28, 6 and 4 targets, respectively. Common axes facilitate comparison of both between-target variation and length-associated changes across methods.

#### Supplementary Fig. S4

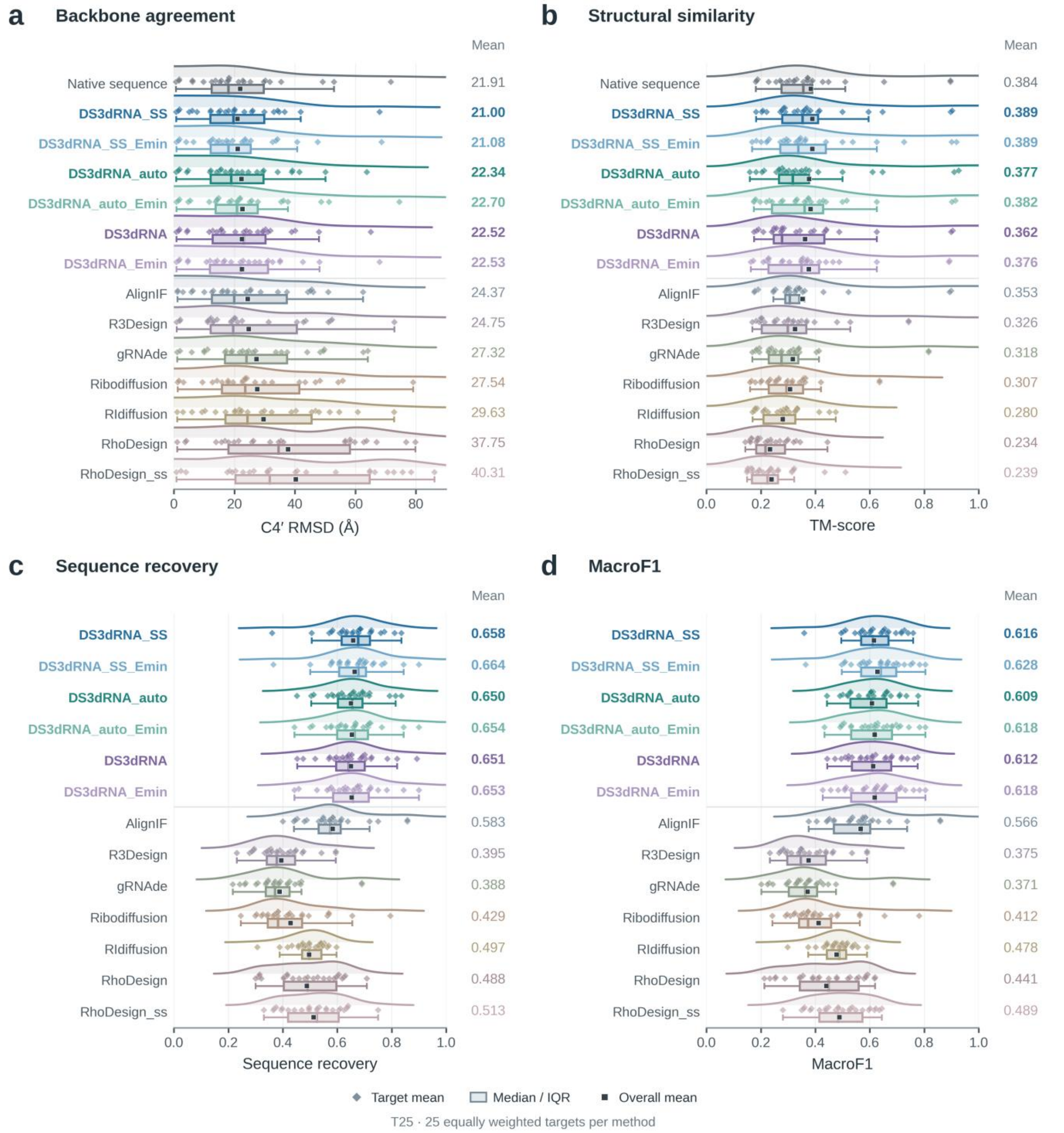

**Supplementary Fig. S4 | Comparison of all DS3dRNA configurations and reference methods on T25.** a,b, C4′ root-mean-square deviation (RMSD; a) and TM-score (b) between AlphaFold3[12] predictions and their experimental target structures. Each target-level structural value is the arithmetic mean across five predicted models. Native-sequence predictions provide a reference for predictor performance. c,d, Target-level native-sequence recovery (c) and MacroF1 (d) for the design methods. DS3dRNA uses the three-dimensional scaffold alone; DS3dRNA_auto additionally applies internally inferred base-pairing constraints; DS3dRNA_SS uses externally supplied secondary structure. The suffix _Emin denotes the additional candidate selected by minimum TriRNASP-Fine energy, assessed separately from the corresponding consensus candidate. Sequence metrics for _Emin entries describe the selected candidate; ordinary DS3dRNA sequence metrics summarize retained per-run outputs. In all panels, diamonds represent the 25 targets; boxes span the 25th–75th percentiles, internal lines mark medians, and whiskers extend to the most extreme values within 1.5 times the interquartile range. All target observations are shown. Squares and numerical annotations indicate arithmetic means across equally weighted targets. Curves are boundary-reflected Gaussian kernel density estimates, scaled to a common display height within each row; their heights are not quantitative comparisons of density between methods.

#### Supplementary Fig. S5

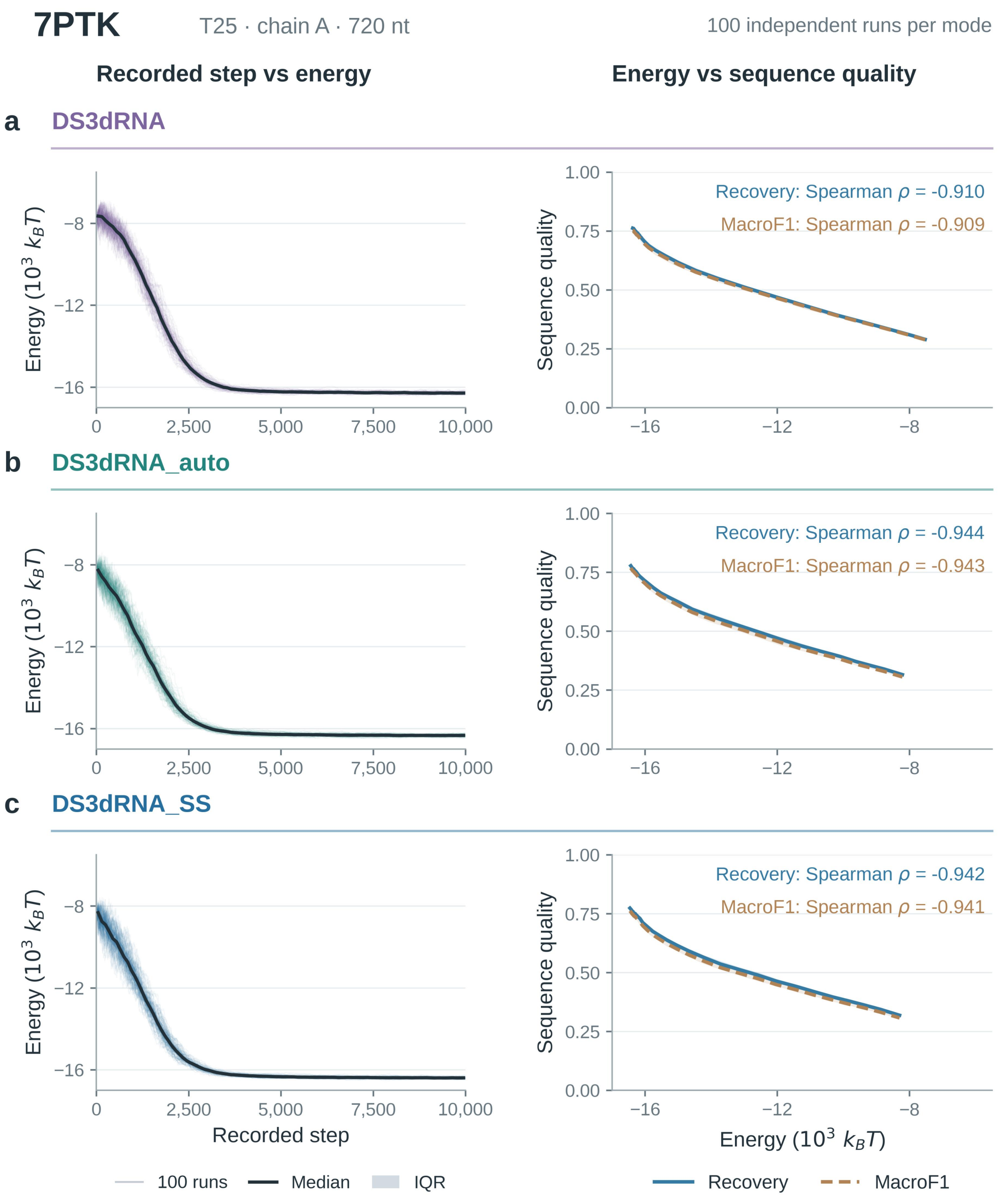

**Supplementary Fig. S5 | Recorded energy profiles and energy–sequence-quality relationships for 7PTK.** a–c, Results for the 720-nucleotide chain A target in T25 using DS3dRNA (a), DS3dRNA_auto (b) and DS3dRNA_SS (c), with 100 independent design runs per configuration. Left, thin colored lines connect retained minimum-energy sequence records in recorded-step order. The dark line and shaded band show the median and interquartile range (IQR), respectively, across within-run mean energies in consecutive 100-step intervals. Right, sequence recovery (blue solid line) and MacroF1 (ochre dashed line) are summarized in 40 quantile intervals of retained energy for each configuration. Each curve and band represents the median and IQR across within-run bin means, with equal weight given to each contributing run. The energy coordinate is the median retained energy in each bin; empty run–bin combinations are omitted. Identical axes are used across configurations, and energy is expressed in units of $10^3$ $k_BT$. Annotated Spearman coefficients[13] are calculated from unbinned retained records pooled across runs, without visit-count weighting: 307,768 records in a, 227,078 in b and 219,631 in c. The trajTopE_10000 archives retain each unique sequence's minimum-energy occurrence within a run and its associated step; they do not reconstruct every state of the complete trajectory. Bands describe between-run IQRs rather than confidence intervals.

#### Supplementary Fig. S6

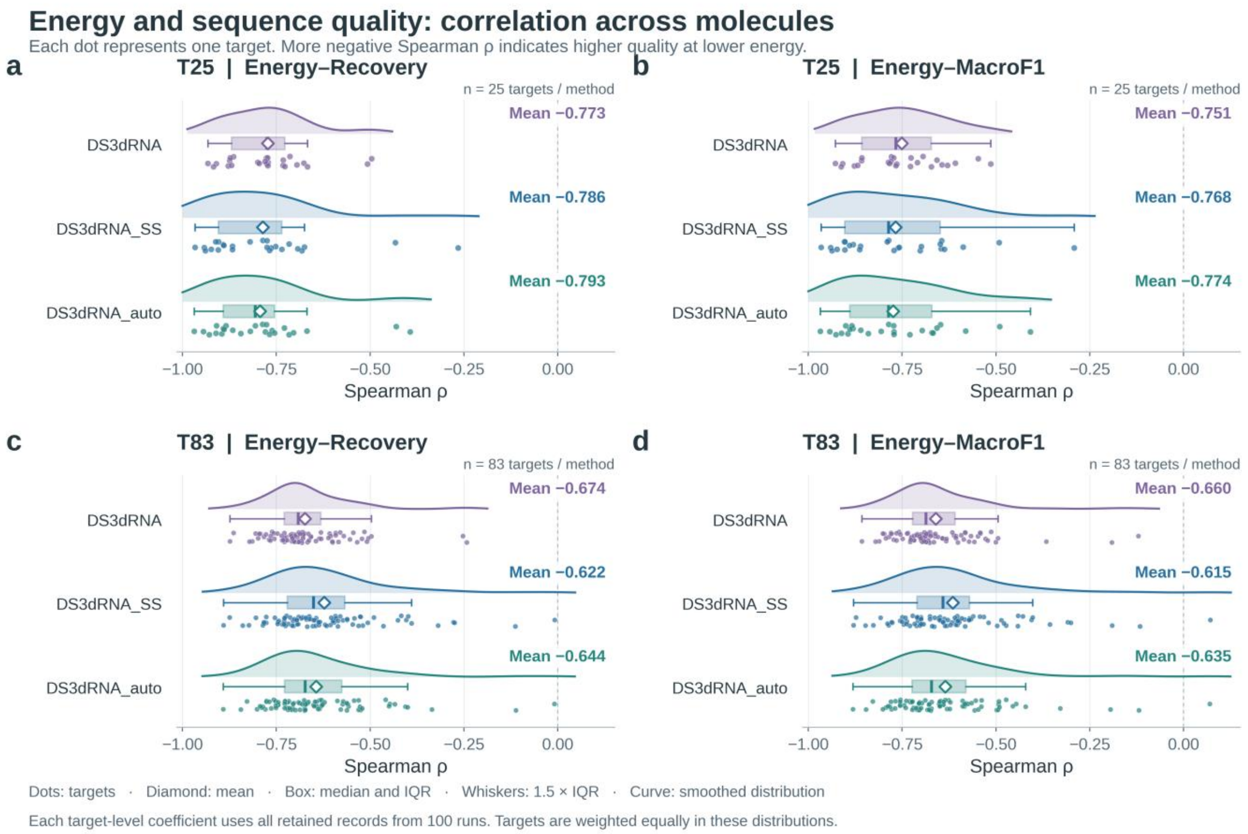


**Supplementary Fig. S6 | Target-level associations between design energy and sequence quality.** a,b, Distributions of energy–recovery (a) and energy–MacroF1 (b) Spearman rank correlations for T25. c,d, Corresponding distributions for T83. Each point is one target-specific coefficient for the indicated DS3dRNA configuration, calculated from all unbinned retained sequence records pooled across 100 independent design runs. T25 and T83 contribute 25 and 83 equally weighted targets per configuration, respectively. Correlations use average ranks for ties and no weighting by sequence visit count. Boxes span the 25th–75th percentiles, internal lines indicate medians, and whiskers extend to the most extreme values within 1.5 times the interquartile range. All coefficients are shown as points, with vertical jitter for visibility. Open diamonds and numerical labels indicate arithmetic means across targets. Curves are descriptive Gaussian kernel density estimates using Scott's bandwidth[14], clipped to the valid correlation range of −1 to 1. Negative coefficients indicate that lower design energy is associated with higher sequence recovery or MacroF1. These summaries describe the retained records; they do not establish independence of records within a run or imply a significance test. Record retention and per-run summaries are described in Supplementary Fig. S5 and Supplementary Note 1.

#### Supplementary Fig. S7

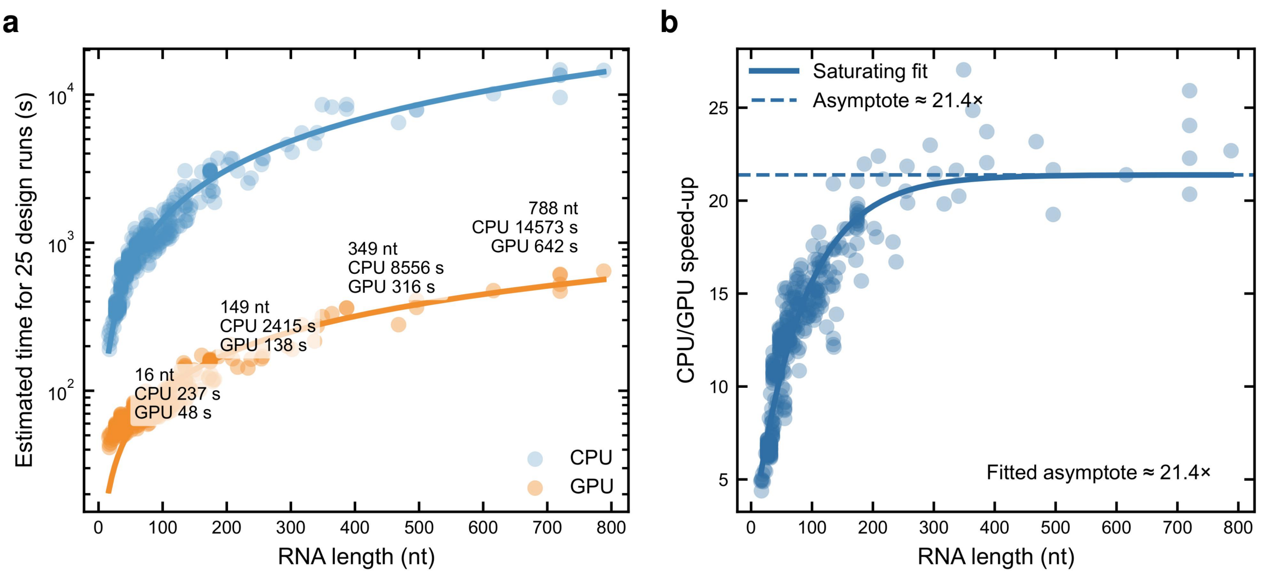


**Supplementary Fig. S7 | Estimated CPU and GPU runtime across RNA lengths.** a, Estimated elapsed times for DS3dRNA jobs configured for 25 design runs of 10,000 Monte Carlo steps, evaluated on a CPU using 16 threads and an NVIDIA GeForce RTX 5080 GPU. Blue and orange points show CPU and GPU estimates, respectively, for the 359 structural inputs spanning 16–788 nucleotides in the supplied runtime workbook. Solid curves are descriptive power-law fits, and the vertical axis is logarithmic. b, Ratios of paired CPU and GPU runtime estimates. The solid curve is a fitted saturating exponential, and the dashed line denotes its fitted asymptote of approximately 21.4-fold. The benchmark script estimated elapsed runtime from short probes by combining the observed probe duration with the progress display's estimated remaining time. The displayed values are runtime estimates, rather than timings of completed jobs. The asymptote summarizes the fitted length dependence and is not an observed maximum speed-up.

#### Supplementary Fig. S8

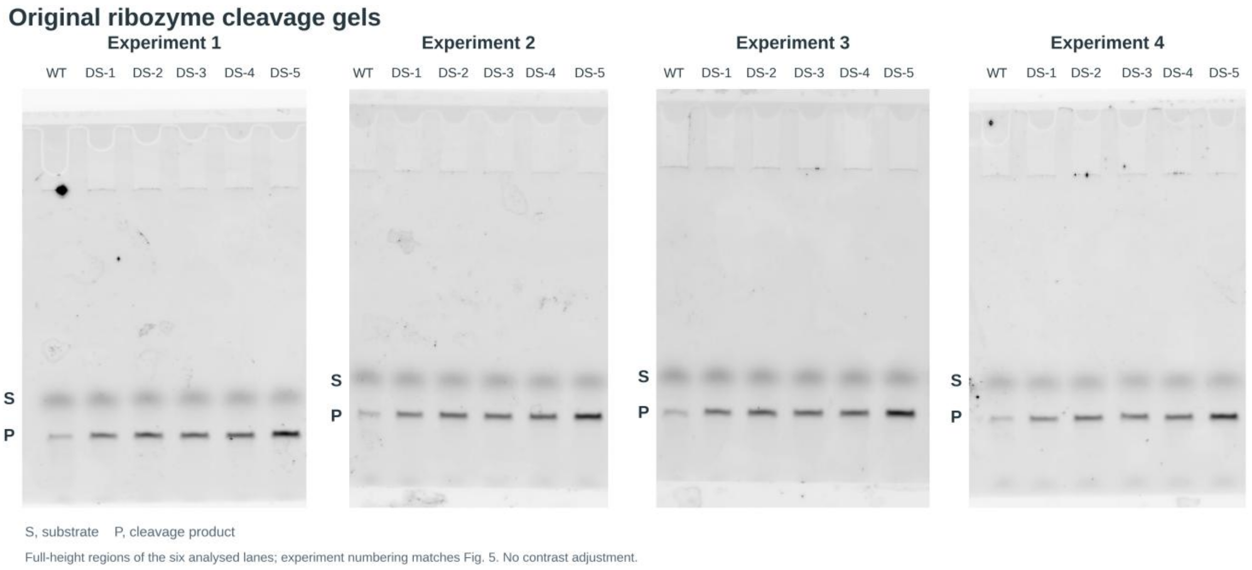


**Supplementary Fig. S8 | Source gel images for the ES2 twister ribozyme cleavage assays.**

Representative source gel images corresponding to the cleavage assays quantified in Fig. 5. Each gel contains the analyzed lanes for wild type (WT) and DS-1–DS-5 designs. S, uncleaved substrate; P, cleavage product. Four independent experiments are shown. Images were cropped from the original scans while preserving the complete analyzed lane regions. No selective band processing was applied.

**Supplementary Table S1 | Raw fluorescence titration measurements for wild-type and designed Mango II aptamers[15,16].** Fluorescence intensities are reported in relative fluorescence units (RFU) for three replicate titration series before normalization. Each of six RNAs (WT, DS-1, DS-2, DS-3, Rh-1 and Rh-2) was measured at 12 RNA concentrations spanning 0–100 nM, with TO1-3PEG-Biotin held at 1 nM, yielding 216 measurements. Excitation and emission wavelengths were 510 and 535 nm, respectively. The zero-RNA measurement in each replicate series provides its normalization baseline. Buffer conditions, normalization and the tight-binding quadratic model[15,17,18] are described in Methods.

| Manuscript sample | RNA (nM) | RFU rep 1 | RFU rep 2 | RFU rep 3 |
|---|---|---|---|---|
| WT | 0.00 | 47 | 63 | 70 |
| WT | 0.05 | 75 | 61 | 67 |
| WT | 0.10 | 83 | 73 | 69 |
| WT | 0.20 | 90 | 75 | 70 |
| WT | 0.50 | 94 | 120 | 118 |
| WT | 1.00 | 140 | 154 | 149 |
| WT | 2.00 | 178 | 193 | 184 |
| WT | 5.00 | 196 | 194 | 193 |
| WT | 10.00 | 205 | 207 | 185 |
| WT | 20.00 | 212 | 220 | 193 |
| WT | 50.00 | 200 | 215 | 218 |
| WT | 100.00 | 214 | 209 | 194 |
| DS-1 | 0.00 | 50 | 43 | 52 |
| DS-1 | 0.05 | 44 | 65 | 56 |
| DS-1 | 0.10 | 62 | 58 | 60 |
| DS-1 | 0.20 | 54 | 75 | 65 |
| DS-1 | 0.50 | 105 | 94 | 100 |
| DS-1 | 1.00 | 121 | 120 | 131 |
| DS-1 | 2.00 | 174 | 150 | 158 |
| DS-1 | 5.00 | 176 | 195 | 160 |
| DS-1 | 10.00 | 200 | 197 | 188 |
| DS-1 | 20.00 | 234 | 196 | 213 |
| DS-1 | 50.00 | 235 | 221 | 225 |
| DS-1 | 100.00 | 217 | 221 | 205 |
| DS-2 | 0.00 | 41 | 58 | 44 |
| DS-2 | 0.05 | 31 | 56 | 30 |
| DS-2 | 0.10 | 46 | 63 | 48 |
| DS-2 | 0.20 | 50 | 67 | 51 |
| DS-2 | 0.50 | 92 | 84 | 52 |
| DS-2 | 1.00 | 123 | 100 | 94 |
| DS-2 | 2.00 | 123 | 122 | 145 |
| DS-2 | 5.00 | 172 | 150 | 148 |
| DS-2 | 10.00 | 175 | 168 | 152 |
| DS-2 | 20.00 | 220 | 171 | 172 |

| Manuscript sample | RNA (nM) | RFU rep 1 | RFU rep 2 | RFU rep 3 |
|---|---|---|---|---|
| DS-2 | 50.00 | 240 | 176 | 189 |
| DS-2 | 100.00 | 226 | 177 | 190 |
| DS-3 | 0.00 | 50 | 46 | 55 |
| DS-3 | 0.05 | 55 | 64 | 59 |
| DS-3 | 0.10 | 59 | 66 | 56 |
| DS-3 | 0.20 | 70 | 76 | 64 |
| DS-3 | 0.50 | 100 | 81 | 104 |
| DS-3 | 1.00 | 135 | 120 | 121 |
| DS-3 | 2.00 | 171 | 164 | 117 |
| DS-3 | 5.00 | 201 | 181 | 148 |
| DS-3 | 10.00 | 214 | 184 | 162 |
| DS-3 | 20.00 | 216 | 195 | 154 |
| DS-3 | 50.00 | 219 | 190 | 163 |
| DS-3 | 100.00 | 221 | 187 | 149 |
| Rh-1 | 0.00 | 32 | 38 | 51 |
| Rh-1 | 0.05 | 55 | 53 | 52 |
| Rh-1 | 0.10 | 36 | 32 | 31 |
| Rh-1 | 0.20 | 26 | 43 | 40 |
| Rh-1 | 0.50 | 37 | 35 | 28 |
| Rh-1 | 1.00 | 59 | 48 | 39 |
| Rh-1 | 2.00 | 47 | 39 | 44 |
| Rh-1 | 5.00 | 33 | 27 | 25 |
| Rh-1 | 10.00 | 40 | 43 | 42 |
| Rh-1 | 20.00 | 55 | 51 | 37 |
| Rh-1 | 50.00 | 29 | 41 | 47 |
| Rh-1 | 100.00 | 44 | 53 | 49 |
| Rh-2 | 0.00 | 32 | 36 | 47 |
| Rh-2 | 0.05 | 44 | 39 | 43 |
| Rh-2 | 0.10 | 43 | 40 | 52 |
| Rh-2 | 0.20 | 39 | 32 | 43 |
| Rh-2 | 0.50 | 43 | 48 | 37 |
| Rh-2 | 1.00 | 41 | 38 | 44 |
| Rh-2 | 2.00 | 38 | 38 | 33 |
| Rh-2 | 5.00 | 25 | 45 | 39 |
| Rh-2 | 10.00 | 32 | 18 | 48 |
| Rh-2 | 20.00 | 37 | 23 | 47 |
| Rh-2 | 50.00 | 29 | 27 | 47 |
| Rh-2 | 100.00 | 41 | 44 | 44 |

#### Supplementary Note 1 | Interpretation of retained design-run records

The recorded profiles and energy–quality analyses use trajTopE_10000 archives for three DS3dRNA configurations on T25 and T83. Each target–configuration combination contains 100 independent runs, giving 324 combinations and 32,400 runs in total. Within a run, a retained unique sequence is represented by its minimum-energy occurrence and the step at which that minimum was recorded. Repeated visits are not expanded according to their count. This retained-record analysis is separate from the count-weighted final-output summaries and consensus construction used for the sequence and structural benchmarks. The resulting profiles therefore describe retained sequence records rather than complete chronological Monte Carlo trajectories.

For each target and configuration, Spearman correlations[13] between energy and sequence recovery or MacroF1 were calculated from all unbinned retained records pooled across the 100 runs, using average ranks for ties. Per-run coefficients and their median and interquartile range provide complementary descriptions of run-to-run consistency. Pooled-record P values are not reported because records within a run are dependent. The distribution summaries in Supplementary Fig. S6 weight each target equally; a more negative coefficient denotes higher sequence quality at lower design energy, but does not establish an association with experimental function.

The complete 324-page collection of target-level profiles is available as  in the DS3dRNA public data archive on Google Drive.

#### Supplementary data and software

Dataset descriptions, download links and the complete design-run profile collection are available in the DS3dRNA public data archive on Google Drive. The three source workbooks for Figs. 2 and 3 are supplied separately as Supplementary Data 1–3, with intact target-level and model-level records. Supplementary Table S1 retains the raw Mango II titration measurements and replicate identities. The repository's Examples directory provides design and sequence-ranking workflows; The Script provides the public consensus and AlphaFold3 input-conversion utility.

#### Supplementary Table S2 | T83 sequence-design benchmark means

Arithmetic means across 83 equally weighted RNA targets; values are fractions on a 0–1 scale. The six DS3dRNA outputs and seven comparator configurations follow Fig. 2 order. Ordinary DS3dRNA target values are count-weighted means of retained per-run outputs; _Emin values score one selected sequence per target. The Fine-energy filter and common decoding rule are specified in Methods. Individual target values, nucleotide lengths and PDB identifiers are provided in Supplementary Data 1.

| Method | Recovery | MacroF1 |
|---|---|---|
| DS3dRNA_SS_Emin | 0.580 | 0.563 |
| DS3dRNA_SS | 0.579 | 0.560 |
| DS3dRNA | 0.578 | 0.559 |
| DS3dRNA_Emin | 0.577 | 0.560 |
| DS3dRNA_auto | 0.574 | 0.553 |
| DS3dRNA_auto_Emin | 0.572 | 0.555 |
| RhoDesign_ss | 0.497 | 0.447 |
| RhoDesign | 0.495 | 0.425 |
| AlignIF | 0.440 | 0.420 |
| RIdiffusion | 0.353 | 0.343 |
| RiboDiffusion | 0.333 | 0.322 |
| R3Design | 0.299 | 0.261 |
| gRNAde | 0.295 | 0.273 |

#### Supplementary Table S3 | T25 sequence and structural benchmark means

Each structural value is first averaged over five AlphaFold3 predictions[12] for a candidate, then averaged across 25 equally weighted targets. Recovery and MacroF1 are averaged across targets, using retained-run means for ordinary DS3dRNA outputs and one candidate per target for _Emin; the native-sequence row has no design-sequence scores. Full target means and model-level observations are retained in Supplementary Data 2 and 3, respectively.

| Method | Recovery | MacroF1 | C4′ RMSD (Å) | TM-score |
|---|---|---|---|---|
| Native sequence | — | — | 21.91 | 0.384 |
| DS3dRNA_SS | 0.658 | 0.616 | 21.00 | 0.389 |
| DS3dRNA_SS_Emin | 0.664 | 0.628 | 21.08 | 0.389 |
| DS3dRNA_auto | 0.650 | 0.609 | 22.34 | 0.377 |
| DS3dRNA | 0.651 | 0.612 | 22.52 | 0.362 |
| RhoDesign_ss | 0.513 | 0.489 | 40.31 | 0.239 |
| RhoDesign | 0.488 | 0.441 | 37.75 | 0.234 |
| AlignIF | 0.583 | 0.566 | 24.37 | 0.353 |
| RIdiffusion | 0.497 | 0.478 | 29.63 | 0.280 |
| RiboDiffusion | 0.429 | 0.412 | 27.54 | 0.307 |
| R3Design | 0.395 | 0.375 | 24.75 | 0.326 |
| gRNAde | 0.388 | 0.371 | 27.32 | 0.318 |

#### Supplementary Table S4 | Software and computational resources

Runtime estimates compared execution with 16 CPU threads of AMD Ryzen 9 9950X3D against execution on an NVIDIA GeForce RTX 5080 GPU; design calculations used the RTX 5080, and local AlphaFold3 predictions used NVIDIA A100 GPUs.

| Software / resource | Version | Use |
|---|---|---|
| DS3dRNA / TriRNASP[1] | This work | Sequence sampling and three-body energy scoring |
| PyTorch[19] / CUDA[20] | 2.7.1+cu128 | GPU batch evaluation |
| US-align[21] | 20241108 | Structural superposition, redundancy filtering and TM-score |
| x3DNA-DSSR[22] | 1.9.10-linux | RNA base-pair and contact annotation |
| Gemmi[23] | 0.6.4 | PDBx/mmCIF to PDB conversion |
| CD-HIT[24] | 4.8.1 | Training-set sequence redundancy filtering |
| Parasail[25] | 1.3.4 | Independent sequence identity check |
| Biopython[26] | 1.85 | C4′ RMSD superposition |
| AlphaFold3[12] | 3.0.1 | Five-model structural prediction |
| AlphaFold Server[12] | Web service | Selected reproducibility checks |
| PyMOL[27] | 3.1.0 | Structure visualization |
| RhoDesign[6] | 70b34f28ae | RNA inverse-design comparator; local source revision |
| gRNAde[9] | 2453b18778 | RNA inverse-design comparator; local source revision |
| R3Design[8] | c05dc4b351 | RNA inverse-design comparator; local source revision |
| RiboDiffusion[11] | 3ac7a557f4 | RNA inverse-design comparator; local source revision |
| RIdiffusion[10] | 4029e17ace | RNA inverse-design comparator; local source revision |
| AlignIF[7] | bb9c6a370a | RNA inverse-design comparator; local source revision |
| Python | 3.13.9 | Analysis and figure-generation scripts |
| NumPy | 2.3.4 & 1.26.4 | Numerical arrays, sequence encoding and statistical summaries |
| tqdm | 4.67.3 | Progress reporting during DS3dRNA sampling |
| SciPy | 1.15.3 | Nonlinear curve fitting, Spearman correlations and density estimation |
| pandas | 2.3.0 | Tabular data processing and figure source-data preparation |
| Matplotlib | 3.10.3 | Statistical plots and publication figure panels |
| Pillow | 12.0.0 | Image cropping, resizing and panel composition |
| PyMuPDF | 1.27.2.3 | PDF assembly, rendering and figure validation |
| lxml | 6.0.2 | SVG assembly and vector-figure validation |
| openpyxl | 3.1.5 | Reading and validating Excel source workbooks |
| ViennaRNA | 2.7.2 | Secondary-structure layouts for schematic figure assets |

##### Supplementary Data 1–3 | Source numerical workbooks

Supplementary Data 1 (Supplementary_Data_1_T83_relate_Fig2.xlsx) contains the 83 target rows used in Fig. 2, PDB identifiers, nucleotide lengths, and recovery/MacroF1 for six DS3dRNA and seven comparator configurations. The final "Average" row is a derived summary and is excluded when reading target rows.

Supplementary Data 2 (Supplementary_Data_2_T25_relate_Fig3.xlsx) contains 25 target-level rows for Fig. 3, including five-model mean C4′ RMSD and TM-score, sequence recovery and MacroF1; its Consensus_vs_E_min sheet distinguishes consensus from energy-selected DS3dRNA outputs. The final "Average" row is a derived summary.

Supplementary Data 3

(Supplementary_Data_3_T25_All_mode_relate_Fig3.xlsx) contains the model-level AlphaFold3 structural metrics for all 25 targets and candidate groups, including five prediction models per group. Model_level is the wide representation and Long_model is the analysis-ready long representation. Structural entries represent predictions, not independent experiments.

27. Schrödinger, LLC. The PyMOL Molecular Graphics System, version 3.1.0. https://pymol.org/
